# Integration of Alzheimer’s GWAS, 3D genomics, and single-cell CRISPRi non-coding screen implicates causal variants in a microglial enhancer regulating *TSPAN14*

**DOI:** 10.1101/2025.04.01.646442

**Authors:** Shannon Laub, Natalia Tulina, Matthew Hoffman, Joe Faryean, Sandhya Ramachandran, Khanh Trang, Stephanie Lewkiewicz, Alessandra Chesi

## Abstract

While GWAS have been successful in providing variant-to-trait associations for human complex diseases, functional dissection of the discovered loci has lagged behind. Here, we describe a variant-to-gene (V2G) mapping effort for Alzheimer’s disease (AD) to implicate causal variants and effector genes from the most recent AD GWAS meta-analyses (101 loci). We leveraged our genomics datasets comprising high-resolution promoter Capture C, ATAC-seq, and RNA-seq from brain-relevant cell types to fine-map AD GWAS variants, identifying 89 candidate causal SNPs and 69 effector genes. We then designed a single-cell CRISPRi screen to perturb candidate regulatory regions (n=74) and assess the transcriptional response in the human microglial cell line, HMC3. Our screen across ∼97,000 cells identified 19 regulatory regions and 19 effector genes. We then elected to functionally dissect our top hit, the *TSPAN14* locus, and we show that an intronic region containing AD-associated SNPs rs7080009, rs1870138, and rs1870137 is a microglia-specific enhancer, with the AD risk haplotype increasing its activity. CRISPR precise genomic deletion of this region decreases *TSPAN14* expression, alters specific cellular pathways including cell adhesion, and decreases secreted levels of pro-inflammatory cytokines IL-6 and IL-8, which are known biomarkers of aging and AD. Our work provides a systematic framework to map GWAS signals to their effector genes for AD and other brain-related disorders, and provides robust leads to follow up with in-depth functional investigations.

## INTRODUCTION

Alzheimer’s disease is a complex disorder with a strong genetic component, characterized by the deposition of amyloid-β and hyperphosphorylated tau in the brains of affected individuals. Genetic evidence from early-onset familial forms of the disease, caused by mutations in the amyloid precursor protein (APP) and presenilins — which cleave APP to release amyloid-β peptides — underscores the critical role of the amyloidogenic pathway. However, therapeutics targeting amyloid-β have largely failed to show clinical efficacy ^1^, emphasizing the urgent need to identify novel targets for drug development. Genetic association studies (GWAS) of the more common late-onset form of the disease (LOAD) have identified numerous loci with small effect sizes ^2–5^. GWAS, however, can only identify the location of association signals, but not pinpoint the underlying causal variants or effector genes. Functional dissection of GWAS loci has proven challenging for several reasons: 1) the signals usually reside in non-coding regions of the genome which are poorly annotated; 2) many SNPs in high linkage disequilibrium are associated at a single locus, requiring fine-mapping to uncover the true causal variants; 3) the effector gene is not always the closest gene, because of long-range gene regulation between cis-regulatory elements (CREs) and their targets; and 4) gene regulation is highly cell type-specific, and the brain is a complex tissue composed of many cell types.

To address these challenges, we have developed a cell-type specific variant-to-gene (V2G) mapping approach based on ATAC-seq, high-resolution promoter Capture C, and RNA-seq ^6^. Briefly, for each cell type of interest: ATAC-seq is used to physically fine-map AD-associated SNPs, Capture C to link them to their effector genes, and RNA-seq to test the expression of the candidate genes, all in the same cell type.

Here, we applied this approach to investigate the two largest and most recent Alzheimer’s disease (AD) GWAS findings, encompassing 101 loci from Bellenguez *et al.* ^2^ and Wightman *et al.* ^3^. Using our published human genomic datasets ^7–11^, we analyzed three major brain cell types: microglia, neurons, and astrocytes. This analysis identified 89 candidate regulatory variants and 91 effector genes, including 69 coding and 22 non-coding genes. We went on to validate these results by using an orthogonal approach, a pooled CRISPR interference (CRISPRi) enhancer screen with single-cell RNA-seq readout in a microglial cell model, the human cell line HMC3. Microglia are the resident immune cells of the brain and are believed to play a key role in the pathogenesis of AD by contributing to phagocytosis of the amyloid-β plaques and promoting neuroinflammation ^12^. Notably, cis-regulatory elements of immune cells, including microglia, are enriched in AD genetic signals more than other cell types ^13–15^ and many AD genes are selectively expressed in microglia ^2,16,17^. Our single-cell enhancer screen identified 19 CREs containing AD-associated SNPs and 19 effector genes and is, to our knowledge, the first large-scale functional assessment of AD GWAS signals (i.e., ∼100 targeted regions). Finally, we functionally dissected the top hit from our screen, a region containing three AD SNPs in high LD (rs7080009, rs1870138, and rs1870138) residing in an intron of the *TSPAN14* gene. We provide evidence via luciferase assays and precise CRISPR/Cas9 deletions that this region is a microglia-specific enhancer regulating *TSPAN14* expression and influencing cell adhesion and secretion of neuroinflammatory interleukins.

## RESULTS

### Variant-to-gene mapping in brain-relevant cell types

We have developed a genomic approach based on high-resolution promoter Capture C, ATAC-seq, and RNA-seq to identify candidate causal variants and effector genes from GWAS for human complex traits ^6–8,10,11,18–26^ (**Figure 1a**). Briefly, we identify a list sentinel SNPs from GWAS of a trait of interest, expand it to consider all SNPs in high linkage disequilibrium (r^2^ >0.7), and intersect their positions with cell-type-specific ATAC-seq peaks and Capture C loops. This allows us to identify putative causal SNPs in open chromatin contacting an open gene promoter (putative effector gene, which we also require to be expressed in the cell type of interest). We applied this approach by leveraging GWAS signals reported in the two largest, most recent AD GWAS meta-analyses ^2,3^. We extracted 111 sentinel SNPs from these studies (83 from ^27^; 38 from ^3^; 10 index SNPs were in common). We retained one representative SNP from each of 10 pairs that were in high LD (r^2^>0.7), resulting in 101 lead SNPs under consideration in total. For each sentinel SNP, we obtained all proxy SNPs (r^2^>0.7) using the European panel of the 1000 genomes phase 3 v.5 available from SNiPA ^28^. This LD expansion process yielded 2,835 candidate AD-associated ‘proxy’ SNPs. Then, we leveraged our genomic database comprising high resolution promoter-focused Capture C, ATAC-seq, and RNA-seq from 10 brain-relevant cell types, which we categorized in three main groups: 1) neurons, including iPSC-derived neural progenitor cells and iPSC-derived cortical neurons ^7^ and ES-derived hypothalamic progenitors and neurons ^10^; 2) astrocytes, including primary astrocytes ^10^; 3) microglia, including primary monocytes ^8,9^, iPSC-derived microglia (iMg), and HMC3 cells ^11^ (**Suppl. Data 1**).

**Figure 1.**
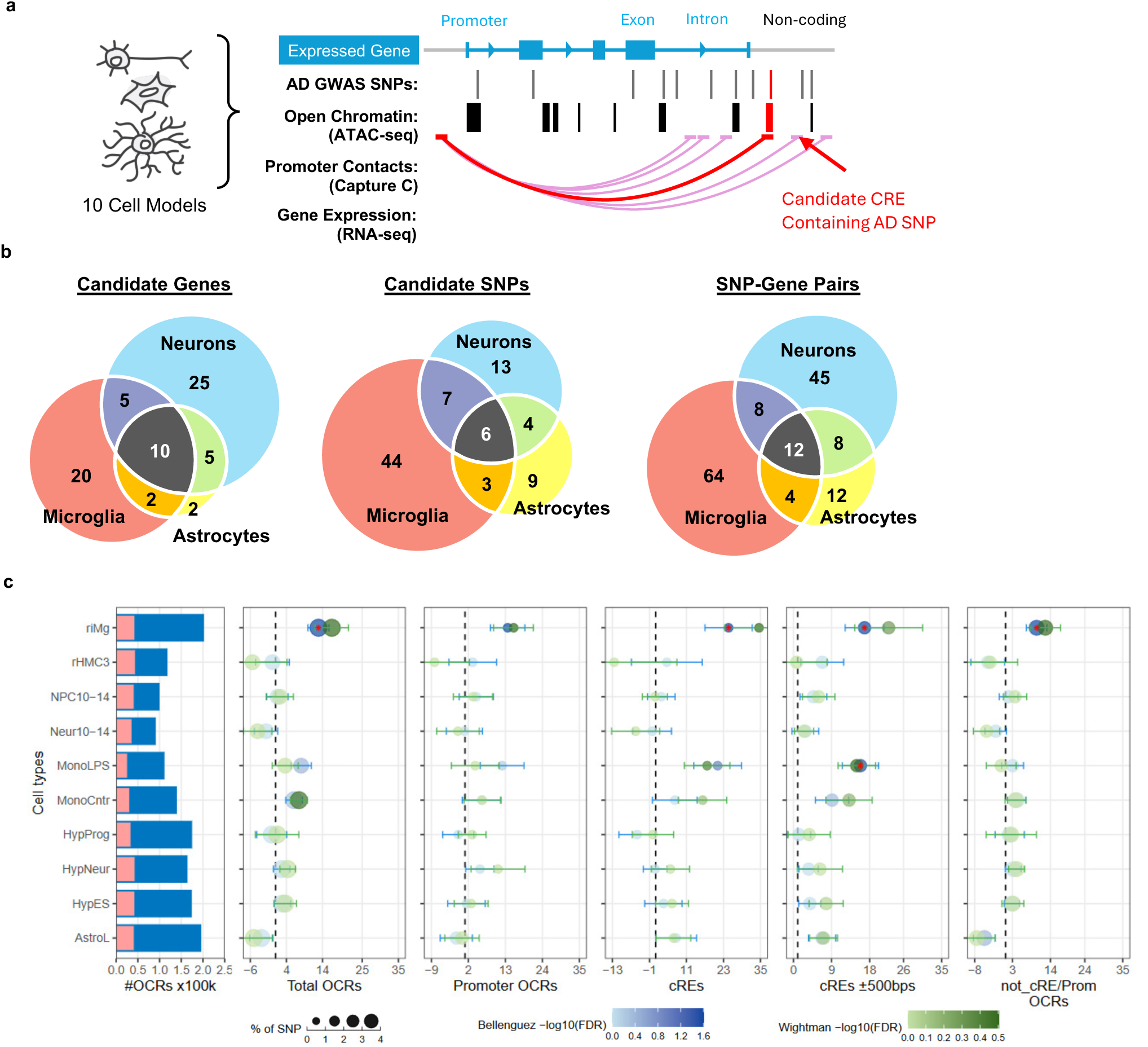
Variant-to-gene mapping identifies candidate causal variants, effector genes, and enriched cell types for AD GWAS loci. **a**: Schematic of the approach used to map GWAS signals to their effector genes. Leveraging our datasets from 10 brain-relevant cell types (5 neuronal, 4 microglial, and 1 astrocyte line; see Suppl. Table 1), we overlaid open chromatin, promoter contacts, and transcriptomics data with AD GWAS SNPs to generate a list of candidate “open” variants contacting “open” actively transcribed genes. **b**: Venn diagrams of the results of V2G mapping showing number of hits and their cell type specificity. **c**: Partitioned LDSC regression analysis showing an enrichment of AD GWAS signals specifically in microglial cell types. Summary statistics from Bellenguez *et al.* and Wightman *et al.* were used for these analyses. Bar-plot shows the total number of open chromatin regions (OCRs) identified by ATAC-seq for each cell type in blue; the proportion of cis-regulatory elements (cREs) identified by Capture-C is shown in pink. The 5 dot-plot panels display the heritability enrichment of AD for each cell type in 5 categories of OCRs: **Total OCRs**: all OCRs identified by ATAC-seq; **Promoter OCRs**: OCRs overlapped with gene promoters; **cREs**: cis-regulatory elements; **cREs ±500bps:** CREs extended by 500 bps**; not_cRE/Prom OCRs**: those OCRs outside of both cREs and promoters. Whiskers are enrichment standard errors. Color-scaled dots correspond to FDR-adjusted *P*-values in -log10, with red asterisks indicating FDR < 0.05. Blue dots show results from Bellenguez *et al*. GWAS and green dots from Wightman *et al*. Dot size corresponds to the proportion of SNP contributing to heritability. Dashed line at 1 indicates no enrichment.

We identified 89 candidate regulatory SNPs in total, of which 44 were specific to microglia, 13 to neurons, and 9 to astrocytes (**Figure 1b**). These variants interacted with 91 promoter regions corresponding to 69 coding genes and 22 non-coding RNAs (**Table 1 and Suppl. Data 2**). Interestingly, we observed different interaction scenarios, including: 1) one enhancer contacting one gene in a specific cell type only (i.e., rs34779859 | rs6733839 contacting *BIN1* in microglial cell types only); 2) one enhancer targeting multiple target genes in a specific cell type only (i.e., rs1377416 targeting *SPI1*, *MIR4487*, and *ACP2* and *NR1H3* in microglial cell types only); 3) one enhancer targeting the same gene in different cell types (i.e., rs74504435 targeting *EGFR* in astrocytes and neurons); 4) multiple enhancers targeting the same gene in one cell type (i.e., *MS4A6A* and *RTF2* in microglia); 5) multiple enhancers targeting the same gene, where the enhancer used depends on the cell type (i.e., *TSPAN14*, *PICALM*, and *FERMT2*). These examples suggest a complex regulation of gene expression where genes can be regulated by different enhancers (which can be cell-type specific or not), and one enhancer can regulate several genes in a given cell type setting.

**Table 1:** Results of the variant-to-gene mapping for AD GWAS in brain-relevant cell types. For each GWAS locus, the sentinel (index) SNP is reported together with the effector SNP rsID, genomic position (in the hg19 build), the r^2^ between the sentinel and the effector SNP, the target gene(s), and the cell group.

| GWAS LOCUS | SENTINEL SNP | EFFECTOR SNP | EFFECTOR SNP LOC | $r^2$ | TARGET GENES | CELL GROUP |
| --- | --- | --- | --- | --- | --- | --- |
| ABCA1 | rs1800978 | rs2777795 | chr9:107672365 | 0.78 | RP11-217B7.3\|TCONS_00016452, SLC44A1 | microglia, astro |
| APP | rs2154481 | rs469420 | chr21:27541944 | 0.71 | ATP5J\|GABPA | neuron |
| BIN1 | rs4663105 | rs34779859 | chr2:127892768 | 0.71 | BIN1 | microglia |
| BIN1 | rs4663105 | rs6733839 | chr2:127892810 | 0.90 | BIN1 | microglia |
| BLNK | rs6584063 | rs117148433 | chr10:98017866 | 0.97 | BLNK, ZNF518A | microglia |
| BLNK | rs6584063 | rs117699776 | chr10:98048265 | 0.87 | BLNK | microglia |
| BLNK | rs6584063 | rs1870169 | chr10:98019157 | 1.00 | BLNK | microglia |
| BLNK | rs6584063 | rs1870171 | chr10:98019113 | 1.00 | BLNK | microglia |
| BLNK | rs6584063 | rs77125544 | chr10:98061083 | 0.84 | ENTPD1-AS1 | neuron |
| CASS4 | rs6069737 | rs544475179 | chr20:55018156 | 0.80 | CASS4 | microglia |
| CASS4 | rs6069737 | rs6024870 | chr20:54997568 | 0.97 | CASS4, RTF2\|TCONS_00028450 | microglia |
| CASS4 | rs6069737 | rs6064392 | chr20:54984768 | 0.91 | CASS4, AURKA\|CSTF1, RTF2\|TCONS_00028450 | microglia |
| CASS4 | rs6069737 | rs718022 | chr20:55003465 | 0.99 | CASS4, AURKA\|CSTF1 | microglia |
| CASS4 | rs6069737 | rs7274581 | chr20:55018260 | 0.87 | CASS4 | microglia |
| CASS4 | rs6069737 | rs76842328 | chr20:55012992 | 0.90 | CASS4 | microglia |
| CASS4 | rs6069737 | rs927174 | chr20:55015166 | 0.89 | CASS4, RTF2\|TCONS_00028450, AURKA\|CSTF1 | microglia, astro |
| CELF1/SPI1 | rs10437655 | rs10838702 | chr11:47410888 | 0.98 | MIR4487, MADD, ACP2\|NR1H3 | neuron |
| CELF1/SPI1 | rs10437655 | rs11039225 | chr11:47430599 | 0.95 | MIR4487, MADD, ACP2\|NR1H3, PACSIN3 | microglia, neuron |
| CELF1/SPI1 | rs10437655 | rs1377416 | chr11:47416746 | 0.97 | MIR4487, SPI1, ACP2\|NR1H3 | microglia |
| CELF1/SPI1 | rs10437655 | rs71475909 | chr11:47377283 | 0.82 | MADD, ACP2\|NR1H3, PACSIN3, ARFGAP2\|PACSIN3, ARFGAP2 | neuron, astro |
| CHRNE | rs7209200 | rs9911764 | chr17:4965589 | 0.94 | SPAG7 | microglia |
| CLU | rs1532278 | rs1532276 | chr8:27466157 | 0.99 | CHRNA2, CCDC25\|ESCO2\|RNU6-1276P, PTK2B, PTK2B\|TRIM35 | neuron |
| CLU | rs1532278 | rs1532277 | chr8:27466181 | 0.99 | CHRNA2, CCDC25\|ESCO2\|RNU6-1276P, PTK2B, PTK2B\|TRIM35 | neuron, microglia |
| CLU | rs1532278 | rs1532278 | chr8:27466315 | 1.00 | PTK2B, PTK2B\|TRIM35 | neuron |
| CLU | rs1532278 | rs2279590 | chr8:27456253 | 0.89 | PTK2B | microglia |
| DOC2A | rs1140239 | rs1140239 | chr16:30021402 | 1.00 | C16orf92\|DOC2A | astro |
| DOC2A | rs1140239 | rs4318227 | chr16:29984839 | 0.77 | HIRIP3\|INO80E, C16orf92\|DOC2A, FAM57B, ASPHD1\|SEZ6L2 | neuron, microglia |
| EPHA1 | rs3935067 | rs3935067 | chr7:143104331 | 1.00 | AC073342.12\|CASP2 | astro |
| FERMT2 | rs7146179 | rs7149638 | chr14:53346931 | 0.80 | FERMT2 | astro, microglia |
| FERMT2 | rs7146179 | rs8003334 | chr14:53374967 | 0.73 | FERMT2 | astro |
| FERMT2 | rs7146179 | rs8019279 | chr14:53374777 | 0.73 | FERMT2 | astro |
| FERMT2 | rs7146179 | rs8019585 | chr14:53374843 | 0.73 | FERMT2 | astro |
| HLA | rs6605556 | rs9271579 | chr6:32590637 | 0.86 | HLA-DRA | microglia |
| HLA | rs6605556 | rs9271580 | chr6:32590641 | 0.87 | HLA-DRA | microglia |
| HLA | rs6605556 | rs9271608 | chr6:32591588 | 0.95 | HLA-DRA | microglia |
| HS3ST5 | rs785129 | rs785143 | chr6:114645237 | 0.90 | HDAC2\|RP3-399L15.3 | neuron |
| KAT8 | rs889555 | rs1549299 | chr16:31154146 | 0.90 | AC135050.5\|ZNF668, STX4 | microglia, neuron |
| KIR3DL2 | rs1761461 | rs1761452 | chr19:54815366 | 0.86 | LILRA5 | microglia |
| KLF16 | rs149080927 | rs113598561 | chr19:1819234 | 0.87 | TCF3, MBD3, MUM1 | neuron |
| MAPT | rs199515 | rs199523 | chr17:44848517 | 0.80 | TCONS_00025674\|WNT3\|WNT9B, WNT3\|WNT9B | astro, neuron, microglia |
| MAPT | rs199515 | rs199524 | chr17:44848438 | 0.80 | TCONS_00025674\|WNT3\|WNT9B, WNT3\|WNT9B | astro, neuron, microglia |
| MAPT | rs199515 | rs70602 | chr17:44859715 | 1.00 | WNT3\|WNT9B, TCONS_00025674\|WNT3\|WNT9B, WNT9B | neuron, astro, microglia |
| MS4A | rs1582763 | rs10736700 | chr11:60031399 | 0.85 | MS4A6A | microglia |
| MS4A | rs1582763 | rs10897011 | chr11:59961427 | 0.76 | MS4A6A | microglia |
| MS4A | rs1582763 | rs11230180 | chr11:59961486 | 0.76 | MS4A6A | microglia |
| MS4A | rs1582763 | rs4938932 | chr11:60033448 | 0.85 | TCONS_00019307, MRPL16, MS4A15 | neuron |
| MS4A | rs1582763 | rs636317 | chr11:60019150 | 0.84 | MS4A6A | microglia |
| MS4A | rs1582763 | rs636341 | chr11:60019161 | 0.84 | MS4A6A | microglia |
| MS4A | rs1582763 | rs672399 | chr11:60020112 | 0.84 | MS4A6A | microglia |
| MS4A | rs1582763 | rs7128450 | chr11:60031270 | 0.85 | MS4A6A | microglia |
| MS4A | rs1582763 | rs718376 | chr11:60002935 | 0.77 | MS4A6A | microglia |
| MS4A | rs1582763 | rs7930318 | chr11:60033371 | 0.85 | TCONS_00019307, MS4A14, MRPL16, MS4A15 | neuron, microglia |
| MS4A | rs1582763 | rs7933202 | chr11:59936926 | 0.74 | MS4A6A | microglia |
| MS4A | rs1582763 | rs7936120 | chr11:60000574 | 0.80 | MS4A6A | microglia |
| MYO15A | rs2242595 | rs2072653 | chr17:18045098 | 0.97 | LLGL1 | neuron |
| MYO15A | rs2242595 | rs2075659 | chr17:18044592 | 0.96 | ALKBH5\|RP11-258F1.1, LLGL1 | neuron |
| NTN5 | rs2452170 | rs2287921 | chr19:49228272 | 0.71 | CA11, CA11\|DBP\|SEC1P | neuron |
| PICALM | rs561655 | rs10792832 | chr11:85867875 | 0.78 | PICALM | microglia |
| PICALM | rs561655 | rs1237999 | chr11:85815030 | 0.92 | PICALM | astro, neuron |
| PICALM | rs561655 | rs7110631 | chr11:85856187 | 0.82 | PICALM | microglia |
| PLEKHA1 | rs7908662 | rs2421016 | chr10:124167512 | 0.99 | PLEKHA1\|TCONS_00018622 | microglia |
| PLEKHA1 | rs7908662 | rs56029211 | chr10:124127990 | 0.98 | PLEKHA1\|TCONS_00018622 | neuron |
| PTK2B | rs73223431 | rs28834970 | chr8:27195121 | 0.99 | PTK2B | microglia |
| PTK2B | rs73223431 | rs73223431 | chr8:27219987 | 1.00 | PTK2B | astro |
| PTK2B | rs73223431 | rs755951 | chr8:27226790 | 0.80 | EPHX2 | microglia |
| RHOH | rs2245466 | rs2245466 | chr4:40198846 | 1.00 | RHOH, TCONS_I2_00020498 | neuron |
| SEC61G | rs76928645 | rs74504435 | chr7:54949256 | 0.95 | EGFR\|EGFR-AS1, EGFR | astro, neuron |
| SHARPIN | rs34173062 | rs34173062 | chr8:145158607 | 1.00 | GPAA1, CTD-3065J16.9\|EXOSC4, FOXH1\|PPP1R16A\|TCONS_00015381 | microglia, neuron |
| SORL1 | rs74685827 | rs74685827 | chr11:121353077 | 1.00 | RP11-730K11.1\|SORL1 | microglia |
| TPCN1 | rs6489896 | rs7297749 | chr12:113659752 | 1.00 | SLC8B1 | microglia |
| TPCN1 | rs6489896 | rs73208737 | chr12:113635055 | 0.84 | SLC8B1 | microglia |
| TPCN1 | rs6489896 | rs76097225 | chr12:113659621 | 0.94 | SLC8B1 | microglia |
| TPCN1 | rs6489896 | rs78810900 | chr12:113679598 | 0.75 | SLC8B1 | microglia |
| TSPAN14 | rs6586028 | rs1870137 | chr10:82269848 | 0.98 | TCONS_00018255 | microglia |
| TSPAN14 | rs6586028 | rs1870138 | chr10:82269611 | 0.98 | TCONS_00018255, TSPAN14 | microglia |
| TSPAN14 | rs6586028 | rs7080009 | chr10:82269461 | 0.98 | TSPAN14 | microglia |
| TSPAN14 | rs6586028 | rs9633740 | chr10:82265271 | 0.99 | TCONS_00018255, TSPAN14 | microglia, neuron |
| TSPOAP1 | rs2632516 | rs2526377 | chr17:56410041 | 0.98 | BZRAP1-AS1, BZRAP1-AS1\|MIR4736 | microglia |
| WDR12 | rs139643391 | rs72934512 | chr2:203926271 | 1.00 | NBEAL1\|WDR12, ICA1L | microglia |
| WOR81 | rs35048651 | rs1310 | chr17:1641715 | 0.94 | AC130689.1\|RPA1\|SMYD4 | astro |
| WOR81 | rs35048651 | rs1317708 | chr17:1639965 | 0.94 | SERPINF2, AC130689.1\|RPA1\|SMYD4 | neuron, astro, microglia |
| WOR81 | rs35048651 | rs2070863 | chr17:1648502 | 0.94 | AC130689.1\|RPA1\|SMYD4 | neuron |
| WOR81 | rs35048651 | rs2287322 | chr17:1641035 | 0.92 | AC130689.1\|RPA1\|SMYD4 | astro, microglia |
| WOR81 | rs35048651 | rs34962442 | chr17:1640529 | 0.93 | AC130689.1\|RPA1\|SMYD4 | microglia |
| WOR81 | rs35048651 | rs4989024 | chr17:1639719 | 0.94 | SERPINF2, AC130689.1\|RPA1\|SMYD4 | neuron, astro, microglia |
| WOR81 | rs35048651 | rs62090051 | chr17:1640640 | 0.93 | AC130689.1\|RPA1\|SMYD4 | microglia |
| WOR81 | rs35048651 | rs8077638 | chr17:1640793 | 0.94 | AC130689.1\|RPA1\|SMYD4 | astro |
| WOR81 | rs35048651 | rs874424 | chr17:1640034 | 0.94 | SERPINF2, AC130689.1\|RPA1\|SMYD4 | neuron, astro, microglia |
| ZCWPW1/<br>NYAP1 | rs7384878 | rs12539172 | chr7:100091795 | 0.76 | AGFG2 | neuron, astro |

Microglia were the most enriched cell type for candidate regulatory SNPs and interactions, suggesting this cell type is the main mediator of AD genetic effects, possibly through microglia specific enhancers.

To investigate this further, we performed Stratified Linkage Disequilibrium Score Regression (S-LDSR) analysis ^29^ across all cis-regulatory elements (CREs) identified through our ATAC-seq and chromatin capture analyses for each cell type utilized in our V2G effort, utilizing the same AD GWAS summary statistics ^3,27^. We assessed three types of genomic regions, i.e. total OCR (all open chromatin regions defined by ATAC-seq), promoter OCRs (the subset of OCRs overlapping a gene promoter), and CREs (the subset of OCRs that do not overlap a gene promoter, but show a chromatin loop to a gene promoter, as assessed by Capture C). As shown in **Figure 1c**, only microglial cell types showed high and significant enrichment for AD GWAS signals, consistently with our V2G mapping findings. Using the Bellenguez *et al*. summary statistics, we found that iPSC-derived microglia were significantly enriched in all categories except “Promoter OCRs” and activated monocytes were enriched in the “CREs ± 500bp” category. Results from the Wightman *et al*. summary statistics were similar, with iPSC-derived microglia significantly enriched in the “Total OCRs” category and activated monocytes in the “CREs ± 500bp” category.

### Single-cell CRISPRi non-coding screen in a microglial cell line

We sought to validate our V2G mapping candidates using an orthogonal approach. As illustrated in **Figure 2a**, we performed a pooled CRISPRi screen at low MOI wherein we targeted the CRISPRi machinery at our candidate CREs and measured for perturbation of gene expression using scRNA-seq. We elected to perform the screen in a tractable microglia cell model, the HMC3 cell line, which is easy to passage and transduce while still recapitulating core microglia markers and characteristics ^30,31^. We designed a library of 227 sgRNA guides targeting 76 candidate regions containing AD associated SNPs from our V2G mapping plus additional SNPs from unpublished data (**Suppl. Data 3**). As positive controls, we designed guides targeting the transcription start site of 7 genes expressed in this cell line (as assessed by our bulk RNA-seq data ^11^), including 3 guides targeting RAB1A that we had previously validated via bulk CRISPRi knockdown experiments. As negative controls, we used 23 non-targeting guides (Sigma Millipore).

**Figure 2.**
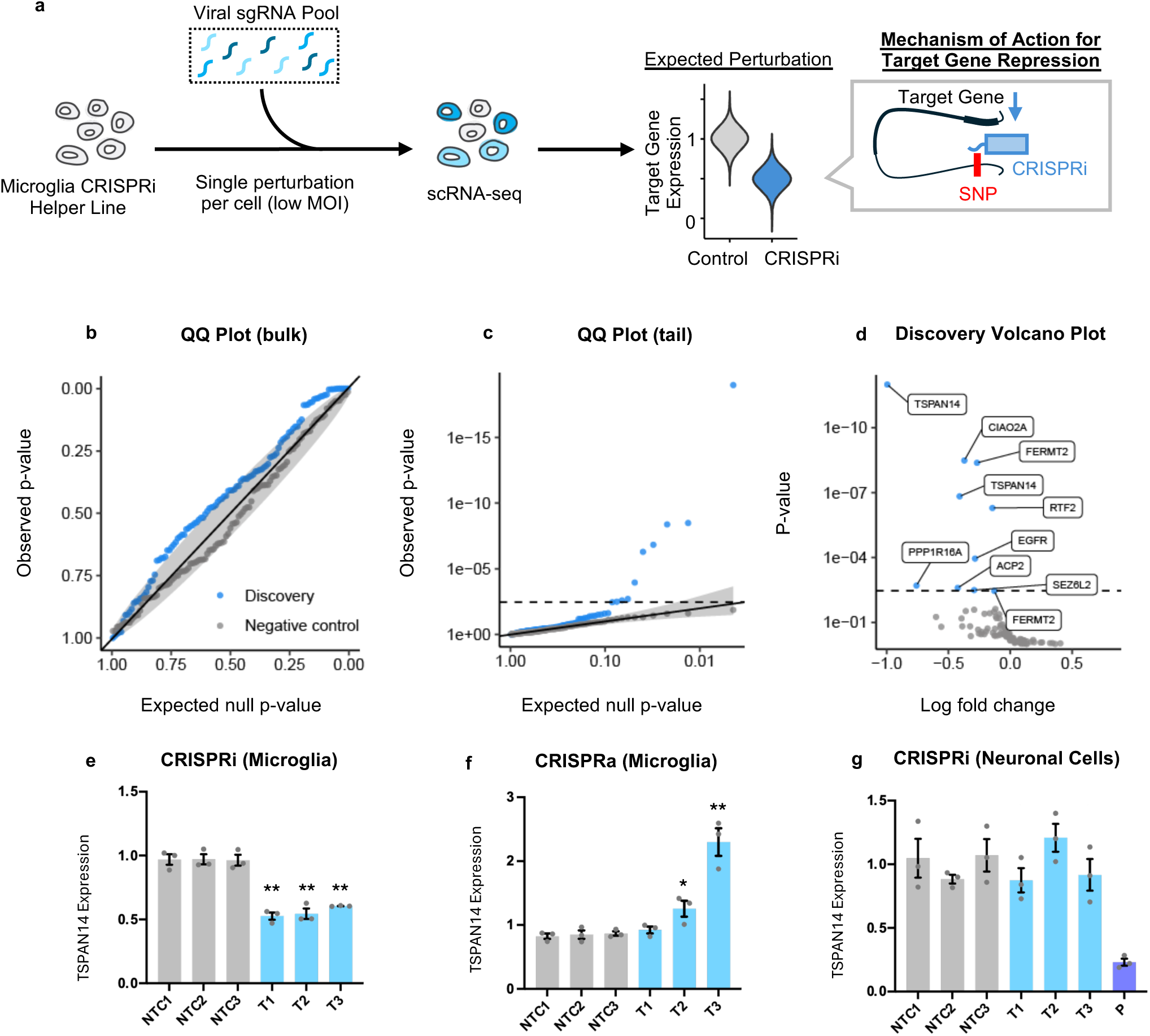
Single-cell CRISPRi screen in microglial cells identifies effector genes for AD GWAS loci. **a:** Schematic of the screen. Three sgRNAs were designed to target each of 74 candidate regulatory regions (CREs) from our V2G mapping effort. We created a CRISPRi helper line expressing dCas9-KRAB in HMC3 cells and transduced it with a lentiviral pool of the sgRNAs at low MOI so that most cells would only receive a single perturbation. After selection and expansion, we performed scRNA-seq with 10x Genomics. Upon successful targeting of a CRE using CRISPRi, we anticipate RNA expression knockdown of the target gene. **b, c, d:** QQ plots from SCEPTRE differential expression analysis showing **b**) good calibration, **c)** significant signal, and **d**) volcano plot of the hits. These plots refer to the V2G “union” analysis. **e**, **f**, **g**: a candidate region containing three AD-associated SNPs (rs7080009, rs1870138, and rs1870137) regulates *TSPAN14* expression in microglial but not in neuronal cells. CRISPRi (**e**) and CRISPRa (**f**) were performed in HMC3 cells stably expressing dCas9-KRAB and dCas9-VPR, respectively, via lentiviral delivery of three gRNAs targeting the region (T1, T2, and T3) and three non-targeting control guides (N1, N2, and N3). *TSPAN14* expression was assessed by qPCR (N=3; bar plots show mean *TSPAN14* expression with SEM error bars normalized to their respective control line with no guide. Significance by one-way ANOVA followed by pairwise t-test. \*\**P*<5×10^−5^; \**P*<0.05. A similar CRISPRi experiment was performed in the neuronal ReNcell VM line, showing no effect on TSPAN14 expression (**c**). P: a guide targeting *TSPAN14* promoter was used as a positive control.

We engineered an HMC3 helper line to constitutively express KRAB-dCas9 and transduced it with a pooled lentiviral library at low multiplicity of infection (MOI <0.1), as verified by flow cytometry (**Suppl. Figure 1**). We leveraged the 10x Genomics Chromium platform, capturing 96,639 cells and sequencing their single-cell transcriptomes and gRNA barcodes on the Illumina platform. After quality control, 74,456 high quality cells were retained for analysis. Each guide was detected in an average of 250 cells with a minimum of 21 and a maximum of 677 cells per guide (**Suppl. Figure 2**). For differential expression analysis, we used SCEPTRE ^32^, which performs analysis of single-cell perturbation screens via conditional resampling.

We first performed an analysis informed by our V2G mapping, testing all our candidate variant-gene pairs, using the following parameters: gRNA-to-cell assignment method: “mixture”; sidedness of test: “left”; and gRNA integration strategy: “union” (**Figure 2b**). This analysis yielded 10 significant pairs corresponding to 9 CREs affecting the expression of 8 genes, i.e. *TSPAN14*, *PPP1R16A*, *ACP2*, *CIAO2A*, *SEZ6L2*, *EGFR*, *FERMT2*, and *RTF2* (after excluding pairs where the candidate CRE was closer than 1kb to the candidate gene’s promoter). We then performed a similar analysis using gRNA integration strategy: “singleton” (where the gRNAs are tested individually) and obtained similar results, with the identification of 2 additional significant targets, i.e. *ASPHD1* and *RPA1*. To increase our discovery power, we also performed both “union” and “singleton” analyses to test for cis effects against all genes included within a 500kb region surrounding the candidate CREs, corresponding to 847 and 2043 pairwise tests after QC, respectively. These analyses yielded 20 significant hits, confirming all the results from the candidate analyses (except for *RPA1*) and identifying 10 additional effector genes, i.e. *LRP4*, *TMEM219*, *PRXL2A*, *DOT1L*, *VKORC1*, *GNB2*, *CYFIP2*, *BRD2*, *CSTF1*, and *EMP3*.

Overall, we found that the “union” method provided more power to detect subtler knockdowns, while the “singleton” analysis was more sensitive when, by chance, one or more guides did not target efficiently (**Suppl. Figure 3**). For all analyses, positive and negative controls demonstrated good calibration (**Suppl. Figure 4**). Non-targeting gRNAs demonstrated no effect, and about half of positive controls demonstrated more than 50% downregulation. Results for all the analyses are reported in **Suppl. Data 4** and **Suppl. Figure 4**, and a summary is shown in **Table 2**.

**Table 2:** Significant CRE-gene pairs identified by our CRISPRi screen in HMC3 cells. All hits from four different SCEPTRE analyses (v2g_union, v2g_singleton, cis_union, and cis_singleton) are compiled here. Most hits were identified in most analyses. Each CRE (cis regulatory element) is identified by the AD-associated SNPs it contains. Relative expression compared to cells containing non-targeting controls comes from the respective SCEPTRE analysis with the greatest knockdown in gene expression.

| **CRE** | **Effector Gene** | **Relative Expression** |
| --- | --- | --- |
| rs7080009\|rs1870138\|rs1870137 | TSPAN14 | 36.46% |
| rs10838702 | ACP2 | 46.22% |
| rs4318227 | SEZ6L2 | 50.26% |
| rs3087616 | CYFIP2 | 50.77% |
| rs9633740 | TSPAN14 | 51.84% |
| rs34173062 | PPP1R16A | 59.10% |
| rs9633740 | PRXL2A | 59.28% |
| rs11039225 | LRP4 | 59.49% |
| rs4318227 | ASPHD1 | 69.97% |
| rs332262\|rs332261 | CIAO2A | 73.42% |
| rs74504435 | EGFR | 74.81% |
| rs8019279\|rs8019585\|rs8003334 | FERMT2 | 76.36% |
| rs6024870 | CSTF1 | 76.94% |
| rs9271608 | BRD2 | 77.50% |
| rs113598561 | DOT1L | 77.94% |
| rs1549299 | VKORC1 | 78.63% |
| rs7149638 | FERMT2 | 80.01% |
| rs2287921 | EMP3 | 84.73% |
| rs12539172 | GNB2 | 85.42% |
| rs34962442\|rs62090051\|rs8077638\|rs2287322 | RPA1 | 85.45% |
| rs927174 | RTF2 | 87.38% |

### Functional dissection of rs7080009, rs1870138, rs1870137 at the *TSPAN14* locus

The top hit from our screen was a region comprising three AD-associated SNPs in high LD (r^2^=1), i.e. rs7080009, rs1870138, and rs1870137, located in an intron of *TSPAN14*. We elected to perform validation and functional dissection of this locus.

We first validated the regulatory effect of this region on *TSPAN14* via bulk CRISPRi and CRISPRa, utilizing the guides from the CRISPRi screen plus additional newly designed guides (**Suppl. Table 1**). To this end, we generated transgenic HMC3 CRISPRi and CRISPRa “helper” lines carrying a lentiviral cassette which expresses the dead Cas9 (dCas9) protein fused to either a transcriptional repressor (KRAB) or activator (VP64) domain. We first transfected them with three “naked” guide RNAs targeting the *TSPAN14* regulatory region and assessed *TSPAN14* expression after 24 hours using qPCR. All three guides targeting the enhancer region significantly decreased *TSPAN14* expression (by 20-60%) in the CRISPRi line, while we did not see an effect in the CRISPRa line (**Suppl. Figure 5**). Given the CRISPRa line with dCas9-VP64 showed no effect, we generated a new CRISPRa line harboring the stronger VPR activating proteins and leveraged lentiviral transductions to constitutively express the guides to avoid problems with the stability of “naked” guides. In this set of experiments, all three targeting guides showed a significant decrease in *TSPAN14* expression (∼50%) in the CRISPRi line, and two out of three guides showed a significant increase (125-225%) in the CRISPRa line (**Figure 2e and f**).

To assess if this regulation was cell-type specific, we performed a similar CRISPRi experiment in the human neural progenitor line ReNcell VM that we have engineered to express the transcriptional repressor ZIM3-KRAB fused to dCas9. We transduced the cells with lentivirus containing the same three sgRNAs targeting the *TSPAN14* enhancer and three non-targeting control guides, and we assessed *TSPAN14* expression by qPCR 96 hours post transduction. Contrary to the results in the HMC3 line, we did not observe an effect in this neuronal line, confirming a microglia specific regulation of *TSPAN14* by this AD-associated enhancer (**Figure 2g**). Overall, these experiments validate the single-cell CRISPRi screen findings and are consistent with the hypothesis that the region harboring rs7080009, rs1870138, and rs1870137 functions as an enhancer for *TSPAN14* specifically in microglial cells.

### The region harboring rs7080009, rs1870138, and rs1870137 functions as a microglia-specific enhancer and the AD risk alleles increase its activity

To support this hypothesis, we investigated the enhancer activity of this region (**Figure 3a**) using dual luciferase reporter assays. We selected a 1.6kb region encompassing the open chromatin region containing rs7080009, rs1870138, and rs1870137 and cloned it into a luciferase reporter vector upstream of a minimal promoter sequence. We transfected HMC3 cells with this construct or a control vector containing the minimal promoter only, together with a reference plasmid used for normalization by transfection efficiency, and performed the luciferase assay 48 hours post-transfection. We found a significant 3.8x increase in reporter activity for the enhancer-containing construct compared to the empty control (*P*=8×10^−4^), which demonstrates that this regulatory region can act as an enhancer in microglial cells (**Figure 3b**). To determine if this enhancer is active in microglia only or also functions in other cell types, we repeated this experiment in a human neuronal line, SH-SY5Y (**Figure 3c**). We did not observe any significant enhancer activity in this line, suggesting that this enhancer is active in microglia specifically.

**Figure 3.**
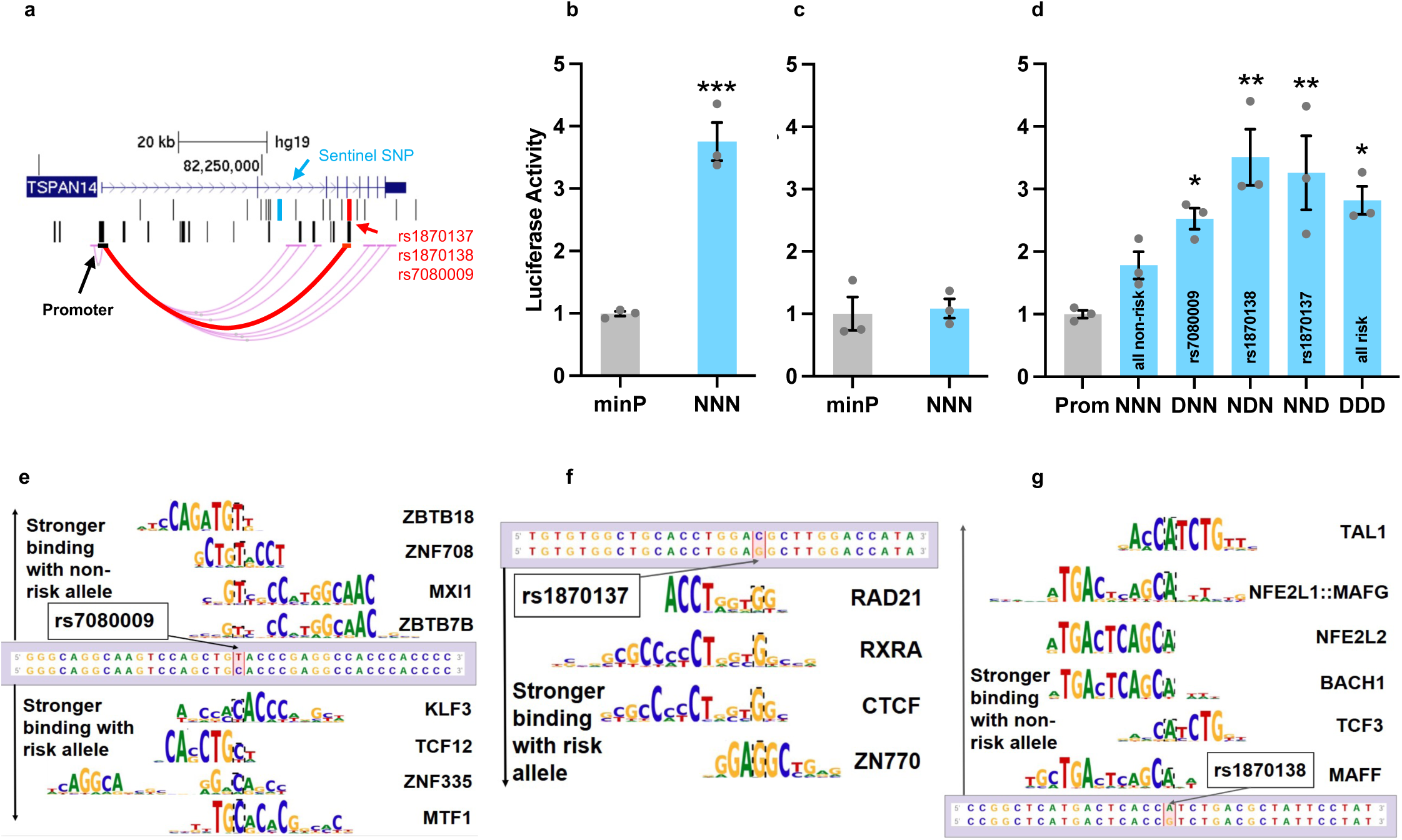
The *TSPAN14* intronic region containing AD-risk variants rs7080009, rs1870138, and rs1870137 acts as a microglia specific enhancer and AD-risk alleles increase enhancer activity. **a:** genomic location of the three AD SNPs (red) showing the chromatin loop with the *TSPAN14* promoter. **b, c**: in luciferase assays, a 1.6 kb intronic region containing the three AD SNPs increases reporter luminescence in microglial cells (**b**; HMC3 cells; two-sample t-test *P*=8×10^−4^), but not in neuronal cells (**c**; SH-SY5Y cells). minP: minimal promoter only (no enhancer region). NNN: enhancer region with 3 SNPs with non-risk alleles. **d**: constructs containing AD risk alleles significantly increase enhancer activity compared to *TSPAN14* promoter alone (* *P*<0.05, ** *P*<0.005), while a construct with all non-risk allele does not. Cell line: HMC3; DNN: rs7080009 risk allele only; NDN: rs1870138 risk allele only; NND: rs1870137 risk allele only; DDD: 3 SNPs with risk alleles; Prom: *TSPAN14* promoter only (no enhancer region). Bar plots show mean reporter activity with SEM error bars normalized to promoter only control. Statistical analysis by one-way ANOVA (*P*=1.7×10^−3^) followed by pairwise t-test with BH correction, N=3. **e, f, g:** rs7080009, rs1870138, and rs1870137 AD-risk alleles affect binding of specific transcription factors in microglial cells. TF binding analysis was performed with motifbreakR using the “default” scoring method. In each panel, sequence logo diagrams for TF binding site motifs matching each SNP locus are plotted with respect to the reference genome (top, non-risk allele; bottom, risk allele). Motifs on top have stronger binding to the non-risk allele and motifs on bottom to the risk allele; both are ordered by the magnitude of the difference in binding affinities. The name of the TF is adjacent to each motif. Motifs depicted have expression > 10 TPM in at least two of the three cell lines analyzed (HMC3, iMg, monocytes).

To assess direction of effect, i.e. whether the AD-risk haplotype increases or decreases *TSPAN14* expression, we first interrogated the GTEx Portal (accessed on 2/5/2025) and found that these SNPs are eQTLs for several genes, including *TSPAN14*. The AD risk alleles were found to correlate with increased *TSPAN14* expression in brain cortex and cultured fibroblasts, while they were correlated with decreased expression in skin, adipose, colon, muscle, esophagus, breast, and nerve tibial tissues.

To test directly the effect of the AD risk SNPs in a microglia cell type, we generated luciferase constructs harboring an enhancer region with combinations of the three alleles at rs7080009, rs1870138, and rs1870137. We cloned them upstream of the endogenous *TSPAN14* promoter and assessed their enhancer activity in HMC3 cells.

Enhancer regions containing one or more AD risk alleles all showed significantly higher activity than the control condition (*TSPAN14* promoter only), while the enhancer containing no AD risk alleles (NNN) did not (**Figure 3d**). These results suggest that AD-risk alleles may increase *TSPAN14* expression in microglia, with a direction of effect consistent with the eQTL data in brain cortex.

#### Transcription factor analysis

A possible mechanism by which SNPs residing in enhancer regions can affect gene expression is by altering binding affinities of transcription factors (TF). Therefore, we performed TF motif analysis at rs7080009, rs1870138, and rs1870137 using motifbreakR ^33^. We included TFs that were expressed (TPM>10) in at least two of our three microglial cell types (HMC3, iMg, and monocytes). We identified 8 TFs (*ZBTB18, ZNF708, ZBTB7B, MXI1, KLF3, TCF12, ZNF335,* and *MTF1*) with the capacity to bind at rs7080009 (**Figure 3e**); the first four showed decreased affinity for the AD risk allele C, while the latter four showed increased affinity for it. We identified 4 TF binding site matches at rs1870137 (*RAD21, RXRA, CTCF*, and *ZN770*), all of which showed increased affinity for the AD risk allele G (**Figure 3f**). Finally, 8 TFs (*TAL1, NFE2L1::MAFG, NFE2L2, BACH1, TCF3, MAFF, NFE2::MAF, and TAL1::TCF3*) had a binding site match at rs1870138, all showing decreased affinity for the risk allele G (**Figure 3g**).

Many of these TFs are known to play an important role in microglia and have been previously associated with AD. Several TFs associated with rs1870138 (*MAF, MAFF, NFE2, BACH1*, and *TAL1*) have been previously identified as master regulators of the microglia gene module, with *TAL1* serving as one consistently across several transcriptome data sets ^34^. *TAL1* is highly expressed in microglia and known to regulate the cell cycle and proliferation in erythroid lineages, a role it is thought to play in microglia, too ^35^. *NFE2L2* activates an antioxidant pathway in glial cells known to protect neurons from death induced by oxidative stress, a hallmark of neurodegeneration in AD ^36–38^. *BACH1* is a transcriptional repressor that negatively regulates *NFE2L2* to modulate brain inflammation ^39^. *NFE2L1* also promotes antioxidant activity in the presence of oxidative stress and regulates proteasome activity, is downregulated in AD, and brain tissue-specific knockout leads to neurodegeneration in mice ^40^.

Among TFs whose binding is altered by rs1870137, *RXRA* is known to regulate development of tissue-specific macrophages ^41^ and cholesterol metabolism ^42^, and variants in the *RXRA* gene itself have been shown to influence AD risk via effects on cholesterol metabolism ^43^. Finally, *MXI1*, whose binding is affected by rs7080009, was shown to be associated with AD pathogenesis in transcriptomic analyses of prefrontal cortical samples of patients with AD ^44^. *MTF1* regulates heavy metals, which are known to be toxic to both neurons and microglia and contribute to AD ^45^. These results provide a list of candidate TFs for further functional characterization in the context of AD.

### Precise genomic deletion of the enhancer encompassing rs7080009, rs1870138 and rs1870137 results in decreased *TSPAN14* expression

To validate the regulatory function of the region including rs7080009, rs1870138, and rs1870137 on *TSPAN14* expression using an orthogonal approach, we generated precise small genomic deletions using CRISPR/Cas9 editing. We designed two guide RNAs flanking the region of interest and cloned them in a lentiviral vector containing Cas9 and one of two selective markers, mCherry or puromycin. We double selected for mCherry (using FACS) and puromycin resistant cells, and derived single cell clones. About 50% of the clones screened by PCR using region-specific primers showed the presence of a genomic deletion. We were able to obtain two homozygous clones (# 1 and # 2, carrying a ∼780 bp homozygous deletion encompassing all three SNPs; **Figure 4a and b**). Deletions were confirmed by Sanger sequencing.

**Figure 4.**
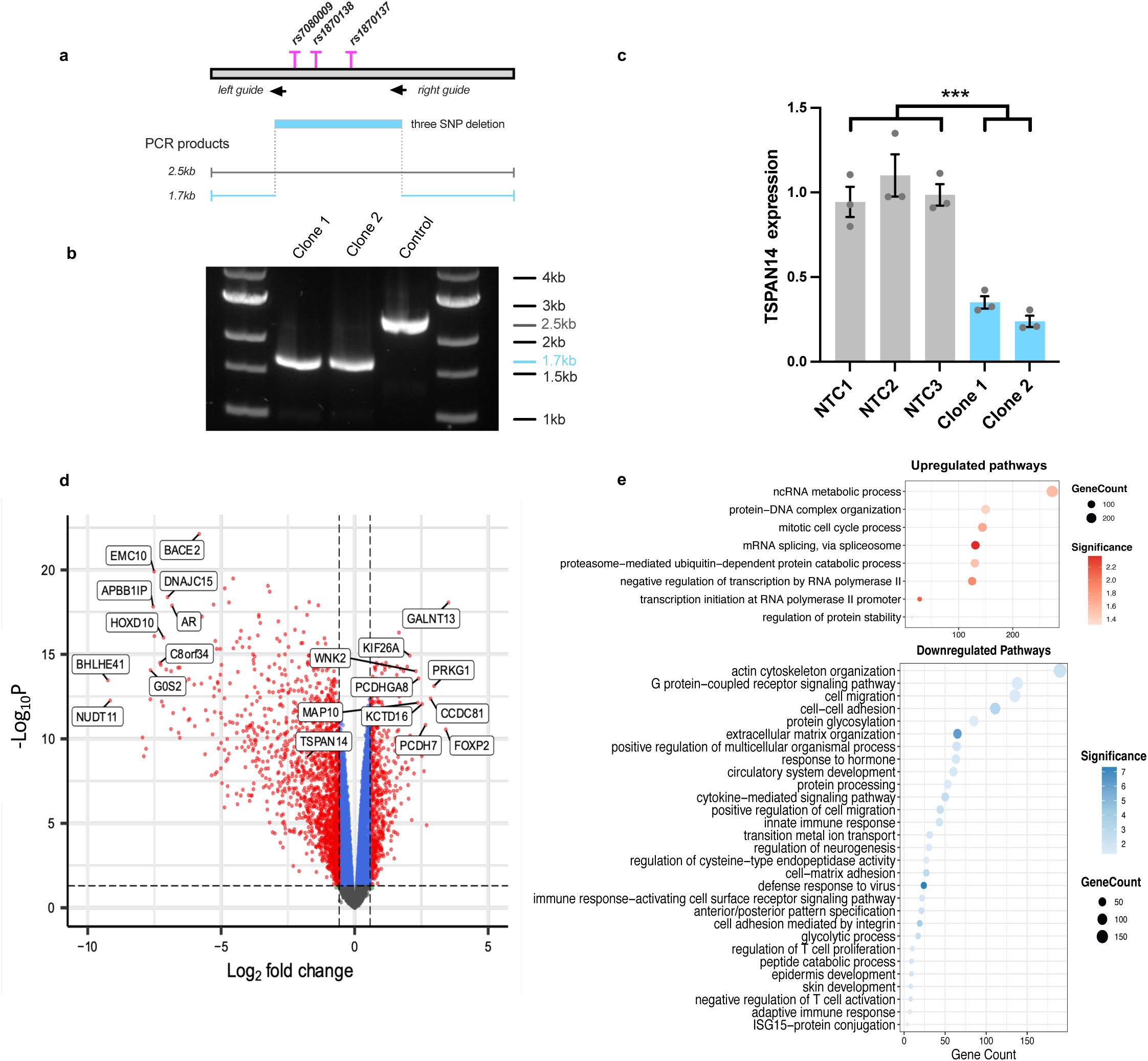
Precise genomic deletions of the *TSPAN14* intronic region containing AD-associated SNPs rs7080009, rs1870138, and rs1870137 downregulate *TSPAN14* expression and cell adhesion and immune response pathways in microglial cells. **a:** via CRISPR-Cas9 editing, we generated two clonal HMC3 lines harboring ∼780 bp homozygous genomic deletions spanning the three AD SNPs. **b:** DNA agarose gel shows sizes of the PCR amplicons for clones 1 and 2 and a control clone with no deletion. **c:** *TSPAN14* expression levels by qPCR in clones 1 and 2 (blue) and control lines (grey). *TSPAN14* expression was significantly decreased (60-80%) in both deletion clones (two-sample t-test *P*=2×10^−3^). NTC1, NTC2, NTC3: control lines, each with a different non-targeting guide. N=3. Bar plots show mean *TSPAN14* expression levels with SEM error bars normalized to the average of the control lines. **d:** Volcano plot showing differentially expressed genes in the same two KO lines vs three control lines expressing non-targeting guides. The dashed lines show thresholds for Fold Change (1.5) and FDR (0.05). **e:** PANTHER enrichment analysis using the GO-Slim Biological Process annotation set. Top (red): significantly upregulated pathways. Bottom (blue): significantly downregulated pathways. Correction: FDR<0.05. For each significantly enriched term, only the top term of the GO hierarchy (i.e., the most specific) is shown.

We then assessed expression levels of *TSPAN14* by qPCR, confirming a strong decrease (∼60-80%) in both lines (*P*<5×10^−3^; **Figure 4c**). These results demonstrate that this AD-associated region is required to maintain a high level of *TSPAN14* expression, consistent with its proposed role as an enhancer.

#### Precise deletion of TSPAN14 enhancer results in significant transcriptomic changes, including downregulation of cell adhesion and immune response pathways

To identify transcriptional changes resulting from deleting the enhancer region, we performed bulk RNA sequencing of the two homozygous enhancer-deleted (KO) clonal lines described above, three Mock lines, and one empty vector transduction control (EV). Each Mock line was transduced with one of three different non-targeting sgRNAs. Three technical replicate libraries were prepared for each line and sequenced on the Illumina platform. PCA and heatmap clustering showed the presence of batch effects due to the libraries being processed in three different batches (**Suppl. Figure 6a and c**). We therefore adjusted for batch effects (see Methods and **Suppl. Figure 6**), which resulted in all technical replicates clustering together in PCA and heatmap plots, with a clear separation by condition (i.e. Mock/EV vs KO, **Suppl. Figure 6b and d**). We then performed differential gene expression analysis between the KO and Mock conditions using a model including batch effects, which yielded 2,287 differentially expressed genes (1,651 upregulated and 636 downregulated). (**Figure 4d** and **Suppl. Data 5**). *TSPAN14* was significantly downregulated in the enhancer KO lines, consistent with our qPCR results (FC = 0.29; FDR = 1.3 × 10^−9^). Pathway enrichment analysis using PANTHER ^46^ with the GO-Slim Biological Process annotation set indicated 29 downregulated and 8 upregulated pathways. Many of the top downregulated pathways were related to cell adhesion (i.e. ‘extracellular matrix organization’, ‘cell adhesion mediated by integrin’, ‘cell-cell adhesion’) and the immune response (i.e. ‘defense response to virus’, ‘cytokine-mediated signaling pathway’, ‘innate immune response’), while top upregulated pathways included mRNA splicing, transcription, and cell cycle (**Figure 4e**).

#### Precise deletion of TSPAN14 enhancer reduces levels of cell adhesion

Since several pathways related to cell adhesion were downregulated according to our pathway analysis, we elected to investigate defects in cell adhesion in the *TSPAN14* enhancer KO lines using phalloidin staining. Phalloidin specifically binds to filamentous actin (F-actin), a critical component of the cytoskeleton which plays a key role in maintaining cell shape, enabling cell motility, and facilitating cell-cell and cell-extracellular matrix adhesions ^47,48^. Therefore, phalloidin staining allows visualization and quantification of the actin cytoskeleton within cells, which is crucial for assessing how well cells adhere to a substrate. We performed the phalloidin staining assay for three technical replicates of each of the KO lines (clone 1 and 2) and three control lines (NTC1, NTC2, and NTC3). Cells were fixed, stained, and imaged on an automated Zeiss widefield microscope. Average cytoplasmic fluorescence intensity of phalloidin for all the single cells in a well was quantified using CellProfiler ^49^ and R. Both enhancer knockout lines experienced a significant ∼30% decrease in average phalloidin staining compared to controls (**Figure 5c and d**; two-samples t-test; *P*=2.3×10^−3^), consistent with the hypothesis that *TSPAN14* expression is required for cell adhesion.

**Figure 5.**
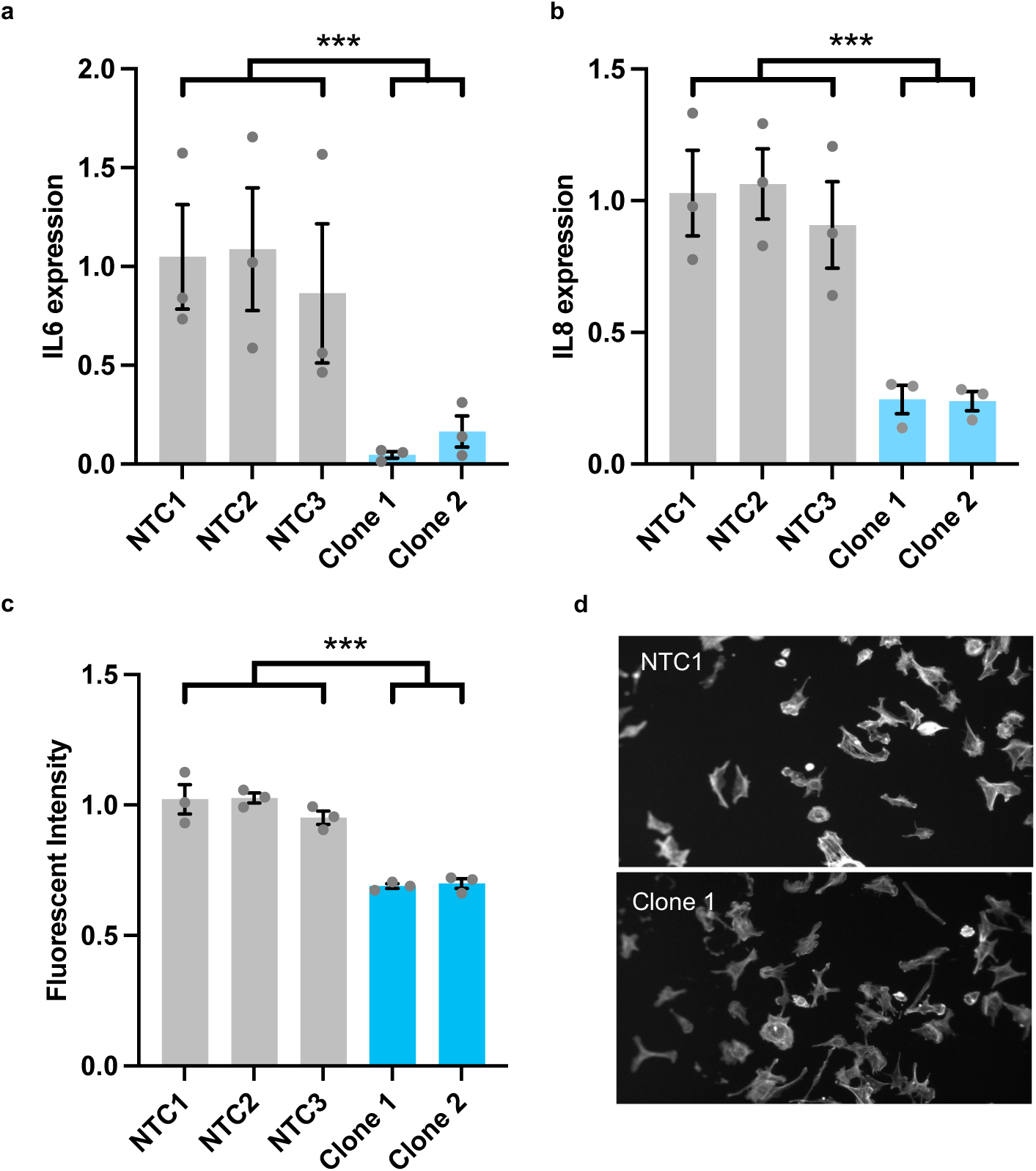
Functional consequences of *TSPAN14* enhancer deletion. **a, b:** Secreted IL-6 and IL-8 protein levels are strongly reduced in *TSPAN14* enhancer KO microglial cells with deletions encompassing AD-risk variants rs7080009, rs1870138, and rs1870137. ELISA was performed on cell culture media of two HMC3 clonal lines (clone 1 and 2, blue). IL-6 or IL-8 protein levels are normalized to the amounts of total protein in each well. Controls are lines harboring a non-targeting guide and/or an empty vector (grey). Bar plots show means with SEM error bars; N=3. **\*\*\*** *P*<5×10^−3^; two-sample t-test. **c**: The same KO lines with *TSPAN14* enhancer deletions (blue) showed a significant reduction (∼25-30%) in intensity of phalloidin (F-actin) staining as compared to three control lines (grey). Single cells were segmented, quantified, and the fluorescence intensities of all cells in each well were averaged. Three wells were measured per condition, with 200-500 cells quantified per well. Bar plots show average fluorescence intensity with SEM error bars. **\*\*\*** *P*<5×10^−3^; two-sample t-test. **d:** Representative images of NTC1 and clone 1 are shown, with notably less F-actin staining observed for clone 1, which harbors the *TSPAN14* enhancer deletion.

#### Precise deletion of TSPAN14 enhancer reduces levels of pro-inflammatory IL-6 and IL-8

Since inflammation mediated by microglia is believed to play a central role in the pathogenesis of AD and cytokine signaling pathways were downregulated in our transcriptomic analysis, we surveyed expression changes in genes linked to inflammation in our *TSPAN14* enhancer KO lines. We noticed that several interleukins and interleukins receptors were downregulated (**Suppl. Data 5**), including the pro-inflammatory cytokines IL-6, which is a biomarker of AD and correlates with cognitive decline ^50–52^, and IL-8, whose levels are also increased with age and in Alzheimer’s disease ^53^. To investigate differences in secreted IL-6 and IL-8 levels, we performed ELISA on the media from our two enhancer KO and three control lines. Secreted IL-6 levels of both IL-6 and IL-8 were strongly reduced in the two KO lines (IL-6: ∼90% reduction; *P*=3×10^−3^; IL-8: ∼75% reduction; *P*=1×10^−3^; **Figure 5 a and b**).

## DISCUSSION

In this study we present a comprehensive variant-to-gene mapping effort for the two most recent Alzheimer’s disease GWAS meta-analyses ^2,3^. Starting from 101 GWAS signals, we created a list of 89 candidate causal SNPs and 91 effector genes leveraging our previously generated genomic datasets (ATAC-seq, high resolution promoter-focused Capture C, and RNA-seq) from 10 brain-relevant cell types. While other studies have performed similar mapping efforts for AD using 3D epigenomics ^54–56^, our work is unique in several aspects: 1) we utilized a wide breadth of cell lines (10 datasets) from three major brain cell types, i.e. neurons, astrocytes and microglia; 2) our comprehensive Capture C library included all the promoters of coding genes and non-coding RNAs, including alternative promoters, and did not rely on antibody precipitation of a specific epigenetic marker; 3) we did not utilize post-mortem tissue, which can lead to biased results or loss of sensitivity to detect early pathogenic events.

We went on to validate and expand this set of candidate variant-gene pairs using an orthogonal technique, a pooled CRISPRi screen where we targeted 74 candidate regulatory regions (harboring the AD-associated SNPs) with the transcriptional repressor KRAB and assessed the effects on the single-cell transcriptomes in the human microglial cell line HMC3. This screen yielded 21 hits, 15 of which predicted by our variant-to-gene mapping effort and 6 novel, resulting in 19 effector genes identified. To our knowledge, this is the first large scale CRISPRi screen performed for AD GWAS variants.

We further characterized the top hit, a region containing three AD-associated SNPs in high linkage disequilibrium—rs7080009, rs1870138, and rs1870137—located within an intron of *TSPAN14* and influencing its expression.

Since a possible mechanism by which the SNPs affect disease is via altering binding of transcription factors (TFs), therefore affecting gene transcription, we performed a TF binding analysis for these three SNPs, identifying several TFs with a putative role in AD pathogenesis.

We next showed that 1) the region harboring these three AD-associated SNPs acts as a microglia-specific enhancer with the AD risk allele increasing its activity, and 2) precise CRISPR/Cas9 mediated deletion of this region in clonal lines decreases *TSPAN14* expression levels. Therefore, our study identifies one (or more) of the three variants rs7080009, rs1870138, and rs1870137 as causal variants at the *TSPAN14* GWAS locus, and *TSPAN14* as the effector gene. In contrast, a recent study by Yang *et al*.^55^ reported rs7922621 (in high LD with the three variants we investigated, but more than 18 kb away) as the causal variant at this locus. This, together with our luciferase results which show an effect of each of the three SNPs on enhancer activity, suggests the possibility of multiple causal SNPs operating at the same GWAS locus. Indeed, this scenario is recently emerging as more common than previously thought ^57^.

Finally, we functionally characterized the effects of a precise genomic deletion of the *TSPAN14* enhancer region. Transcriptomic analysis and functional follow up experiments revealed that loss of TSPAN14 expression results in a downregulation of cell adhesion and neuro-inflammatory pathways. Specifically, KO cells showed a decrease in the levels of IL-6 and IL-8 (proinflammatory cytokines linked to aging and cognitive decline, including in AD ^50–53^ both at the transcriptional and at the protein levels.

*TSPAN14* is an understudied member of the tetraspanin family, a group of 33 small transmembrane proteins involved in various cellular processes such as cell proliferation, cell cycle, invasion, migration, apoptosis, autophagy, tissue differentiation, and immune response ^58^, and is highly expressed in microglia. TSPAN14 physically interacts with ADAM10, the main α-secretase that cleaves APP in the non-amyloidogenic pathway, increasing its expression at the cell surface and promoting its maturation ^59,60^. ADAM10 also cleaves TREM2, a key player in microglial biology and AD, releasing soluble TREM2 in the extracellular space ^61^. For these reasons, *TSPAN14* represents a novel, promising target for further functional dissection in the AD context.

In summary, this study integrated several orthogonal epigenomic techniques (Capture C, ATAC-seq, partitioned LDSC regression, TF analysis, and CRISPRi non-coding screens) in brain-related datasets, providing a list of 19 AD effector genes for further functional investigation. By genetic manipulation and functional dissection in a tractable microglial cell model we uncovered a role of *TSPAN14* and its AD-associated enhancer in cell adhesion pathways and interleukin signaling. The direction of effect from our luciferase experiments suggests that decreased TSPAN14 expression linked to the minor alleles at rs7080009, rs1870138, and rs1870137 may protect from AD through reduced cell adhesion and decreased pro-inflammatory cytokines secretion in microglia.

We anticipate that similar studies in other cell types such as neurons, astrocytes, and oligodendrocytes will further advance the functional characterization of AD GWAS variants and provide insights into cell-specific pathogenic mechanisms.

## METHODS

### V2G mapping

#### AD loci analyzed

We extracted all sentinel SNPs for the significant loci reported in the recent AD GWAS meta-analyses from Bellenguez et al.^2^ and Wightman et al.^3^, for a total of 111 SNPs (83 from Bellenguez; 38 from Wightman; 10 index SNPs were in common). We retained one representative SNP from the 10 pairs that were in high LD (r^2^>0.7), resulting in 101 SNPs under consideration in total. We obtained all proxy SNPs in high LD (r^2^>0.7) with these sentinel SNPs using SNiPA ^28^, selecting the European panel of the 1000 Genomes phase 3 v.5 database.

#### Fine-mapping using ATAC-seq

For each brain-relevant cell type in our database, we physically fine-mapped the AD associated proxy SNPs by requiring them to reside in an open chromatin peak from that cell type, generating a list of cell-type specific candidate regulatory variants. We used the union of the open chromatin peaks called with 2 different methods: 1) optimal peaks from the ENCODE ATAC-seq pipeline (https://www.encodeproject.org/atac-seq/) and 2) reproducible peaks called with our own pipeline, where a peak is called if it’s present in the majority of the technical replicates available for a cell type.

#### Gene target identification using high-resolution promoter Capture C

For each cell type, we intersected the coordinates of the cell-type specific candidate regulatory SNPs with high-resolution promoter Capture C data from our database and selected the ones that showed a chromatin loop with a gene promoter. We further required the promoter to be in open chromatin and the corresponding gene to be expressed in the specific cell type under investigation, using cell-specific ATAC-seq and RNA-seq data from our database. The result of this analysis was a list of candidate causal SNP-gene pairs for each cell type.

### Partitioned heritability LD score regression enrichment analysis

#### Definition of cis-Regulatory Elements (cREs)

To ensure consistency across datasets, we lifted all ATAC-seq open chromatin regions (OCR) coordinates from GRCh37/hg19 to GRCh38/hg38. Promoter Capture C loops at both 1-fragment and 4-fragment resolutions were similarly merged and lifted from GRCh37/hg19 to GRCh38/hg38. We then intersected the ATAC-seq OCRs of each cell type with the corresponding chromatin interaction loops and promoters (defined as −1,500 to +500 bp around the transcription start site) from GENCODE v40. A gene promoter is considered open if it overlaps with an OCR. A cRE is defined as an OCR that connects with at least one open gene promoter via a chromatin contact loop.

#### Reformatting of the GWAS summary statistics

we used GWAS summary statistics from Bellenguez *et al*.^2^ and Wightman *et al*.^3^. We lifted the Wightman summary stats (excluding the 23andMe cohort) from GRCh37/hg19 to GRCh38/hg38. The baseline model LD scores, plink files, allele frequencies, HapMap3 variants list and regression weight files for the European 1000 genomes project phase 3 in GRCh38 were downloaded from https://alkesgroup.broadinstitute.org/LDSCORE/GRCh38/.

#### Cell type specific partitioned heritability of each trait

We used LDSC v.1.0.1 with the --h2 flag to estimate the SNP-based heritability of each trait within 5 sets of input regions from each cell type: (1) OCRs, (2) OCRs at gene promoters, (3) cREs, (4) cREs with an expanded window of ±500 bp, (5) OCRs that were not cREs and not at a gene promoter. Each set of input regions from each cell type was used to create the annotation, which in turn was used to compute annotation-specific LD scores for each cell type region of interest. These annotation-specific LD scores were used with 53 categories of the full baseline model (v2.2) to compute partitioned heritability. Partitioned LD scores were compared to baseline LD scores to measure enrichment fold change and enrichment p-values, which were adjusted with FDR across all comparisons.

### Cell Lines and Culture

Human microglial clone 3 (HMC3; ATCC, CRL-3304), HEK293T (ATCC, CRL-1573), and SH-SY5Y (ATCC, CRL-2266) cells were sourced from ATCC and cultured in Eagle’s Minimum Essential Medium (EMEM, ATCC, cat# 30-2003), Dulbecco’s Modified Eagle’s medium (DMEM, Thermo Fisher Scientific, cat#11995065), and DMEM/F12 (1:1, Thermo Fisher Scientific, cat# 21331020), respectively, containing 10% FBS (Fisher Scientific, cat#MT35010CV), and penicillin-streptomycin (10x, Thermo Fisher Scientific, cat#15140122). ReNcell VM were obtained from Sigma Millipore (SCC08) and cultured in Matrigel (Corning, cat# 354234) coated plates using ReNcell growth media: DMEM/F12/Neurobasal (1:1:2) (made by adding equal volumes of DMEM/F12, Thermo Fisher Scientific, cat# 21331020, and Neurobasal Plus media, Gibco, cat# A3582901), 2.5µg/ml insulin (Sigma, cat# I9278), 1x GlutaMAX supplement (100x, Thermo Fisher Scientific, cat# 35050061), 0.5x MEM non-essential amino acids solution (NEAA, 100x, Invitrogen, cat#11140050), 0.5mM sodium pyruvate (ThermoFisher Scientific, cat# 11360070), 0.5x B27 supplement (50x, Thermo Fisher Scientific, cat# 17504-044), 0.5x N2 supplement (100x; Thermo Fisher Scientific, cat# 17502-048), and 1x penicillin-streptomycin solution (10x; ThermoFisher Scientific, cat#15140122) with 20ng/ml of human recombinant factors epidermal growth factor (EGF, R&D systems, cat# 236-EG) and fibroblast growth factor (FGF, R&D systems, cat#233-FB/CF) each. EGF and FGF were added to ReNcell growth media right before use to maintain ReN cells in an undifferentiated state. Cells were maintained at 37 C 95% O2 and 5% CO2. Cell media were changed every 2-3 days and passage when cells reached ∼90% confluency.

### Generation of CRISPRi/a Helper Cell Lines

To generate CRISPRi and CRISPRa helper lines, HMC3s were transduced with lentivirus containing either KRAB-dCas9-T2A-BlasticidinR (Sigma-Aldrich, Cat. CRISPRIE from CRISPRI10X-1KT) or dCas9-VPR-P2A-mCherry (Addgene #154193), respectively. To create the CRISPRi helper line, HMC3s were transduced with CRISPRi lentivirus from the 10X CRISPRi Feature Barcode Optimization Kit (Sigma-Aldrich, Cat. CRISPRI10X-1KT) at a multiplicity of infection (MOI) of 0.1. Cells were incubated with lentivirus for 24 hours before selecting for transduced cells using 7µg/mL blasticidin (Invitrogen, cat# R21001). After 7 days, selection was complete, and surviving cells were maintained under 1µg/mL blasticidin. To create the CRISPRa helper line, HMC3s were transduced with dCas9-VPR-P2A-mCherry that was packaged into lentivirus in-house. Cells were incubated with lentivirus for 24 hours, expanded to a T175 flask, and then sorted for mCherry using flow cytometry.

### Pooled CRISPRi sgRNA Library Design

We designed CRISPRi guides using a custom pipeline. Target regions were first chosen by selecting an interval 99 base pairs upstream and downstream of each SNP of interest (total region size ∼200bp). Overlapping target regions containing SNPs located in closed proximity were merged. The location of each target region was used as an input for CRISPick ^62^ and Flashfry ^63^. The coordinates of this regions were uploaded to CRISPick and a quota of 30 guides were picked using Human GRCh37 as the reference genome, CRISPRi as the mechanism, and SpyoCas9 as the enzyme (using the scoring algorithm from Chen *et al.*) ^64^. CRISPick then generated a ranked list of guides for each region. We then inputted these guides into FlashFry to check for significant off-targets and filter for good GC content (>25% and <75%) and no poly T sequences (more than four sequential Ts in any guide).

After our initial filtering, we noticed that the top ranked guides often had significant sequence overlap (i.e. shifted by 1-3 base pairs from each other). We assumed that guides with similar sequences would have similar on- and off-target effects and would not be experimentally distinct, therefore we chose only the top three guides from each region that were not overlapping one another. After picking our initial set of guides, one region (rs7149638) was difficult to design guides for, so the picking region for this SNP was expanded to 400bp.

### Selecting 10x Chromium Capture Sequence

To select the optimal capture sequence for the 10X Chromium platform, we utilized the 10X CRISPRi Feature Barcode Optimization Kit following the standard protocol. All four capture sequence configurations were tested in the HMC3 CRISPRi helper line. The CS1 sequence in the stem loop had the best sgRNA capture efficiency as compared to CS2 in the stem loop, or either sequence when located in the 3’ end of the sgRNA, and was therefore utilized for the screen.

### Plasmid and lentiviral production

Millipore Sigma performed cloning, viral packaging, and quality control of the CRISPRi sgRNA library. The sgRNA library was cloned into LV13 U6-gRNA-10X:EF1a-Puro-2a-TagBFP (Millipore Sigma), where the CS1 capture sequence was inserted into the stem loop of the sgRNA. The viral titer of the library was 9.1×10^8^ viral particles per mL as determined by a p24 antigen ELISA titer.

### Lentiviral Packaging

In-house lentiviral packaging was performed for the dCas9-VPR-P2A-mCherry construct. HEK293Ts were co-transfected with dCas9-VPR-P2A-mCherry and packaging vectors pMD2.G and psPAX2 (Addgene #12259 and #12260) using Lipofectamine 3000 (Thermo Fisher Scientific, cat# L3000008). Media was exchanged for fresh media 24 hours after transfection. The media was collected 48 and 72 hours after transfection, spun down at 800xg for 3 minutes, and then supernatant was syringe filtered with a 0.45um filter (Fisher Scientific, cat# 13100107). Filtered virus was either transduced into cells immediately or frozen into aliquots and stored at −80°C.

### scRNA-seq Sample Preparation and Sequencing

Several transduction conditions for low MOI were tested empirically in the final 10cm dish format. Viral infection rate was measured using flow cytometry for TagBFP. Final infection conditions were chosen that yielded a 0.1 MOI (∼10% viral infection rate) upon interpolating from tested conditions. Our HMC3 CRISPRi helper line was transduced at 0.1 MOI such that less than 5% of cells would be transduced with more than one sgRNA after selection, as based on Poisson modeling. 3.75 million cells (250k cells/10cm dish) were transduced at an MOI of 0.1 for a guide coverage of at least 1000x cells transduced per guide. Three additional plates were transduced with the same lentiviral preparation to measure transduction efficiency during flow sorting.

Twenty-four hours after transduction, lentiviral media was exchanged for fresh media. Forty-eight hours after transduction, media was exchanged for puromycin selection media and exchanged again every subsequent 48 hours. Six days after transduction, cells were sorted for BFP+ cells containing an sgRNA. Twelve days after transduction, cells were prepped for 10x single-cell barcoding. HMC3s were rinsed twice with PBS, detached with TrypLE (Life Technolologies, cat# 1260402), strained twice with a Flowmi filter (Sigma-Aldrich, cat# BAH136800040), diluted to 1200 cells/μL, and brought to the Center for Applied Genomics Core. Samples were run on the 10x Chromium for single-cell barcoding and Illumina’s NovaSeq for sequencing using the protocol CG000316 from 10x Genomics for Chromium Next GEM single cell 3’ reagent kit v3.1 (dual index) with feature barcode technology to capture the sgRNA sequences for CRISPR screening. The pooled sample was run across twelve 10x Chromium lanes and captured a total of 96,639 single cells.

### Differential Gene Expression Analysis

From the FASTQ files, Cell Ranger v7.1 was used to perform alignment of reads, filtering, single-cell barcode counting, and UMI counting. Next, we used SCEPTRE to assign guides to cells using a mixture assignment method ^32^. High-quality cells were then identified by filtering out any cells containing more than one guide, cells with a high percentage of mitochondrial reads, and any cells with very high or very low number of UMIs sequenced or number of genes expressed in the cell (**Suppl. Figure 2**). We then performed four different types of analyses to identify CRISPRi perturbations. Analyses were performed either with a “union” of all cells containing any of the three guides for a single target or in “singleton” where cells were only grouped by their individual guide. To identify relevant genes that may have been perturbed, gRNA-gene testing pairs were created in one of two ways: 1) based on our Capture C loops that linked specific genes with specific candidate regulatory regions or 2) by pairing candidate regulatory regions to any genes that were located within 500kb of the region of interest. Any gene closer than 1kb to the target region was excluded since the CRISPRi machinery is known to suppress genes within that distance ^65^. We then used SCEPTRE to test these pairs for differential gene expression as compared to NTC containing cells. For all SCEPTRE analyses, we specified low MOI, left-sided testing, using the NTC as our control group, and resampling using permutations. We used the number of nonzero cells and the number of UMIs as our covariates. Next, NTC cells were randomly paired with target genes and tested to measure the false discovery rate. Finally, a discovery analysis was performed to evaluate which genes per a given target site were differentially expressed.

### Transcription Factor Binding analysis

The motifbreakR R package was used to identify transcription factors with a binding site motif match at a SNP and score the comparative strengths of the match in the presence of the major (AD risk) and minor alleles. For each of the three SNPs of interest (rs7080009, rs1870137, and rs1870138), motifbreakR was run with the weighted sum (method = “default”) with parameters: filterp = TRUE, threshold = 1e-04. The scoring methods quantify the strength of the match between the candidate binding site and the TF motif, and the p-value associated with the score serves as a cutoff below which matches are deemed sufficiently strong. The full suite of available human TF motifs (pwmList = human) was tested, consisting of those in the “MotifDb” R package and the supplementary motifbreakR datasets “encodemotif,” “homer,” “hocomoco,” and “factorbook.” After filtering by p-value, TFs with RNA expression levels exceeding 10 TPM in at least one of the three relevant cell types (HMC3, induced microglia, and monocytes) were retained, and binding affinities in the presence of the major and minor allele were compared and noted.

### Generation of single cell lines with deletions in TSPAN14 intron removing rs7080009, rs1870138 and rs1870137

Guide RNAs were designed to target the region encompassing rs7080009, rs1870138 and rs1870137: right guide - ATCAAAACATGGCGGGGATT and left guide - GCAACCACAAGCCGGCAAAG. These guides were cloned in lentiviral vectors lentiCRISPRv2puro (Addgene, #98290) and lentiCRISPRv2mCherry (Addgene, #99154), respectively by using two synthetic oligonucleotides per guide with added BsmBI cut sites (**Suppl. Table 1**) and the Golden Gate Assembly cloning protocol. Both vectors also encode Cas9 (Csn1) endonuclease from *Streptococcus pyogenes* which is expressed upon integrating in the human genome. Viral particles containing lentiCRISPRv2puro-right guide and lentiCRISPRv2mCherry-left guide constructs were generated by co-transfecting 625ng of either vector and helper plasmids pMD2.G (Addgene, #12295) and psPax2 (Addgene, #12260) each (2.5µg of DNA total) in HEK293T cell line (ATCC, CRL-3216, plated 300-500,000 cells per 6-well a day before transfection) using Lipofectamine 3000 reagent (Thermo Fisher Scientific, cat# L3000001). 2ml of cell culture media were collected from each 6 well on the second day after transfection, filtered using 0.45um syringe filters (Fisher Scientific, cat# 13100107) and applied to a 60% confluent 10cm plate with HMC3 cells in the presence of 8µg/ml polybrene (Fisher Scientific, cat# TR1003G) as following: 2ml of lentiCRISPRv2puro-right guide and lentiCRISPRv2mCherry-left guide virus-containing media were mixed with 8 ml of cell culture media for HMC3 cells (EMEM, ATCC, cat# 30-2003, with 10% FBS Fisher Scientific, cat# MT35010CV) and 8µl of 10mg/ml MilliporeSigma polybrene transfection reagent (Fisher Scientific, cat# TR1003G). Transduced HMC3 cells were expanded to three confluent 10cm plates and FACS sorted based on the presence of mCherry signal (FACSJazz, Children’s Hospital of Philadelphia, flow sorting facility). Top 5% of mCherry-positive cells with the highest level of mCherry expression were selected and, after limited expansion, subjected to puromycin selection (5µg/ml) for 10 days. Single puromycin-resistant, mCherry-positive cells were expanded and sorted into several 96 well plates. one cell per plate. Thirty-nine single cell clones were analyzed using PCR on genomic DNA using forward (TGTACCTCGTGACATAGC) and reverse (CATCTGGTGTGGCTACAAAG) primers surrounding the region with the three SNPs. Twenty of these clones produced shorter than the full size (2.5kb) bands indicating the presence of a deletion in the region of interest. Two clones, #1 and #2, with single 1.7kb bands were further analyzed using Sanger sequencing (primer - TGCAGTTCTGTGGCACCATC), which revealed two almost identical alleles in both clones, containing 775bp and 783bp deletions which removed all three SNPs. The allele with the 783bp deletion also contains a short, 15bp, insertion, near the sequence of the left RNA2 guide. In addition, a total of four control cell lines expressing three non-targeting control (NTC) guides cloned in the lentiCRISPRv2mCherry vector and a fourth line with the vector sequence alone (“empty vector” or EV) were obtained using the same Golden Gate cloning method. crRNA sequences for the NTC control lines are:

NTC1 – CTGCATGGGGCGCGAATCA

NTC2 – CGCTTCCGCGGCCCGTTCAA

NTC3 – ATCGTTTCCGCTTAACGGCG.

Corresponding synthetic DNA oligonucleotide sequences which were used for cloning these non-targeting guide RNAs in the lentiCRISPRv2mCherry (Addgene, cat# 99154) vector using Golden Gate Cloning are shown in **Suppl. Table 1**.

### RNA extraction

80% confluent HMC3 cells were lifted from 6 well plates (about 300-400,000 cells per well, one well per sample) using TrypLE Express reagent (Life Technolologies, cat# 1260402) and spun at 130rpm for 5 min. to collect cells. Total RNA extraction was done using QIAshredder columns (QIAgen, cat# 79654) and RNeasy Plus micro kit (QIAgen, cat# 74034) following the manufacturer’s instructions. RNAs were eluted in 30μl of nuclease-free water and their concentrations were determined using Qubit RNA broad range (Thermofisher Scientific, cat# Q10211) and high sensitivity (Thermofisher Scientific, cat# Q32852) kits.

### cDNA synthesis

cDNA synthesis was done using Applied Biosystems High-capacity cDNA reverse transcription kit (Life Technology, cat# 4368814) following the manufacturer’s instructions. The same RNA amount (700ng-1μg) was used as template for each 20μl reaction.

### Genomic DNA purification

Genomic DNA (gDNA) was purified from HMC3 cells carrying TSPAN14 enhancer deletion clones ## 1 and 2 as well as NTC and EV control lines (see Generation of single cell lines with deletions in TSPAN14 intron removing rs7080009, rs1870138 and rs1870137) using DNeasy Blood and Tissue Genomic DNA purification kit (QIAgen, cat# 69504) according to manufacturer’s instructions. gDNAs were eluted in 30μl of nuclease-free water. DNA concentrations were determined using Nanodrop and Qubit double-stranded DNA broad range kit (ThermoFisher Scientific, cat# Q32850).

### PCR genotyping

To identify single cell clones with genomic deletions removing rs7080009, rs1870138 and rs1870137 variants, gDNAs were isolated from candidate enhancer deletion lines as well as NTC and EV control lines (see Genomic DNA purification) and used as templates to amplify the *TSPAN14* enhancer region using forward (TGTACCTCGTGACATAGC) and reverse (CATCTGGTGTGGCTACAAAG) primers (amplicon size - 2,473bp). The reaction was performed using 100ng of gDNA per sample in a 25μl of total volume with Q5 high-fidelity DNA polymerase (New England Biolabs, cat# M0491) in the presence of betaine enhancer (Sigma-Aldrich, cat# B0300, final concentration 1M). PCR amplification was done as following: 1. 94°C, 2min. 2. 94°C, 30sec; 61°C 30sec; 72°C, 75sec (30 cycles), 3. 72°C 5min, 4. 4°C, and the resulting PCR products were analyzed using DNA electrophoresis in a 1% agarose/1xTAE gel. All control samples produced single bands with expected mobility (around 2.5kb) for the full size unedited fragment. In contrast, two single cell clones, ##1 and 2, produced single shorter fragments (around 1.7kb), indicating the presence of deletions within the *TSPAN14* enhancer. These bands were excised from the gel using DNeasy Blood and Tissue Genomic DNA purification kit (QIAgen, cat# 69504) and Sanger sequenced.

### Quantitative PCR

qPCR reactions were performed using PowerTrack SYBR Green Master mix (Thermofisher Scientific, cat# A460120) with primers for *TSPAN14* (forward - GAAGAGCAAGTGGGATGAGTC, reverse - TGCCAGGAAGATGCCAAA; located between exons 8 and 9) and *GAPDH* (forward – TGTAGTTGAGGTCAATGAAGGG, reverse – ACATCGCTCAGACACCATG; located between exons 2 and 3) genes in 10µl of a total volume. Both primer sets recognize all isoforms of the corresponding genes. qPCR reactions were done using QuantStudio 12K Flex (Applied Biosystems) instrument. The data were analyzed using the ΔCT method.

### Guide RNA transfection in HMC3 CRISPRi helper line

HMC3 CRISPRi helper line was obtained by transducing HMC3 cells with KRAB-dCas9-T2A-BlasticidinR lentivirus following a standard protocol. Three guide RNAs targeting the *TSPAN14* enhancer region with rs7080009, rs1870138 and rs1870137 were designed using the online guide RNA design tool CRISPOR ^66^ and selected based on highest ranking: crRNA1 - CAGATCGATATCGTCCCGGTAGG, crRNA 2 - ACAGATACGTAGGCACTCGATGG, crRNA 3 – ACGGTATGCAGCGCCTAAGGAGG. 37.5pmol of each sgRNA were obtained from Synthego and used to reverse-transfect 100,000 of detached CRISPRi HMC3 cells in a 6 well format using Lipofectamine 3000 reagent (Thermofisher Scientific, cat# L3000001). Cell pellets were collected 24 hours after transfection for RNA extraction and qPCR analyses to determine changes in *TSPAN14* expression.

### HiFi cloning

The HiFi cloning approach (NEBuilder HiFi DNA Assembly Master Mix, cat# E2621S) was implemented to clone a PCR amplified *TSPAN14* enhancer region encompassing rs7080009, rs1870138 and rs1870137 (hg19 coordinates: chr10:82,268,673-82,270,263, 1,591bp) in two different plasmids which would be used for dual luciferase reporter assays: pNL[Nlucp/minP/hygro] (Promega; custom-made) containing a minimal promoter (minP) and the Nanoluc reporter cassette (the pNL-minP plasmid), and pGL4.10[luc2] (Promega) containing the endogenous promoter of the *TSPAN14* gene, which was digested out of a commercially available plasmid (Genecopoeia, cat# HPRM34820-PF02) using restriction endonucleases EcoRI (NEB, cat# R3101S) and SacI (NEB, cat# R3156S) (the resulting plasmid: pGL4.10-TSPAN14promoter), and the firefly luciferase reporter cassette (the resulting plasmid: pGL4.10-TSPAN14promoter/enhancer). The following forward and reverse primers were used for cloning *TSPAN14* enhancer and promoter regions in the pNL-minP plasmid:

CCGGTACCTGAGCTCGCTAG*GGAGGGTCCAGGTGGAAC* (forward),

CGCCGAGGCCAGATCTTGATG*CTGAGACCAGGCAAGCC* (reverse) (*TSPAN14* intron sequence is shown in italics; these primers contain BmtI and EcoRV restriction sites); in the pGL4.10-TSPAN14promoter plasmid:

*CCCGCCTAAGCTTGGTACC*GATCAAGATCTGGCCTCG (forward),

*CGCATGAGCCACGGTGCC*ATCCTCGAGGCTAGCGA (reverse), and in the pGL4.10-TSPAN14promoter/enhancer plasmid:

GCCTAACTGGCCGGTACCTGCCGGTACCTGAGCTCGCTAG (forward),

CACGGTGCCTGGCGAATTGCTGAGACCAGGCAAGCC (reverse)

The sequences of all cloned regions were confirmed using Sanger sequencing with a primer GTGGGTTGTCTGCCTGCAGCTT or whole plasmid sequencing (Plasmidsaurus).

Sequencing results showed that the cloned *TSPAN14* enhancer region contains all AD-risk alleles (disease - D) for rs7080009, rs1870138, and rs1870137. These alleles were subsequently replaced with non-risk variants (N) using a tri-partite HiFi cloning system which resulted in a series of pNL-minP and pGL4.10-TSPAN14promoter/enhancer plasmids with different allelic combinations of the three SNPs, specifically, DDD, NDN, DNN, NND and NNN (all five alleles were generated for the pGL4.10-TSPAN14promoter/enhancer plasmid only). The primers used to create these alleles were:

rs1870137; *G* to *C* (D to N): forward – CTGCACCTGGA**C**GCTTGGAC, reverse - GTCCAAGC**G**TCCAGGTGCAG.

rs1870138; *G* to *A* (D to N): forward – CATGACTCACC**A**TCTGACGCTATTCC, reverse - GGAATAGCGTCAGA**T**GGTGAGTCATG.

rs7080009; *C* to *T* (D to N): forward – CAAGTCCAGCTG**T**ACCCGAG, reverse – CTCGGGT**A**CAGCTGGACTTG. The nucleotides which were substituted to produce non-risk alleles of rs7080009, rs1870138 and rs1870137 are shown in bold.

Other primer pairs used were: for the pNL-minP plasmid - CGACGCTCAAGTCAGAGGTGG (forward), CCACCTCTGACTTGAGCGTCG (reverse) and CAGCTCGCATCAACGTCTAAGG (forward), CCTTAGACGTTGATGCGAGCTG (reverse).

### Dual luciferase reporter assays

To determine if an intronic region of the *TSPAN14* gene containing rs7080009, rs1870138 and rs1870137 shows enhancer activity, dual luciferase reporter assays were performed using two similar systems, Nano-Glo Dual-luciferase reporter (Promega, cat# N1610) and Dual-Glo luciferase reporter (Promega, cat# E2920) kits. These assays rely on the simultaneous expression of two distinct luciferase-encoding plasmids, the reporter and reference, allowing to assess the effect of any genetic element, such as the *TSPAN14* intronic region, on the expression of the reporter luciferase gene which is then normalized to the expression of the second, reference, luciferase gene which does not depend on that genetic element and serves as a proxy for transfection efficiency.

#### Nano-Glo Dual-luciferase reporter assay

The 1.6kb candidate enhancer region from *TSPAN14* intron harboring AD SNPs rs7080009, rs1870138 and rs1870137 was cloned in the pNL[Nlucp/minP/hygro] plasmid (Promega; custom-made) containing a minimal promoter (minP) and the Nanoluc reporter cassette (the pNL-minP plasmid) as described above (see HiFi cloning). The allelic combination with all three non-risk alleles of these SNPs (pNL-minPNNN) was compared to a control pNL plasmid with minP alone (pNL-minP). pNL-minPNNN and pNL-minP plasmids were transfected in both a microglial and neuronal cell line, HMC3 and SHSY5Y, respectively, along with a reference plasmid pGL4.53[luc2/PGK] which encodes a Firefly luciferase.

In the HMC3 experiment, 10,000 cells were plated per well in a 96 well plate format and transfected on the following day with 50ng of either pNL-minPNNN or pNL-minP plasmid, 1ng of the reference plasmid and 149ng of transfection carrier DNA (Promega, #E4881). Lipofectamine LTX (ThermoFisher Scientific, #A12621) was used as a transfection reagent. All samples were plated in triplicate.

In the SH-SY5Y experiment, 20,000 cells were plated per well in a 96 well format and transfected with 160ng of either pNL-minPNNN or pNL-minP plasmid and 40ng of the reference plasmid using Lipofectamine 3000 reagent (ThermoFisher Scientific, cat# L3000001). All samples were plated in triplicate.

The Nano-Glo Dual-luciferase reporter assay (Promega, cat# N1610) was performed on the next day after transfection using the GloMax Discover microplate reader (Promega). The ratios of Nanoluc and Firefly luciferase signals were calculated and compared between experimental pNL-minPNNN/pGL and control pNL-minP/pGL conditions. This experiment was performed in three biological replicates.

#### Dual-Glo luciferase reporter assay

To determine the effects of rs7080009, rs1870138 and rs1870137 SNPs on enhancer activity of the *TSPAN14* intronic region, five different allelic combinations of these SNPs, specifically, NNN, DDD, NDN, NND, DNN, were created and positioned upstream from the endogenous promoter of the *TSPAN14* gene in the pGL4.10[luc2] reporter plasmid (Promega, see HiFi cloning). Each of the resulting pGL4.10-TSPAN14promoter/enhancer plasmids, which encode Firefly luciferase, were then transfected in HMC3 cells, along with a reference plasmid (pRL-TK, Promega), which expresses Renilla luciferase. A pGL4.10 plasmid with *TSPAN14* promoter only (pGL4.10-TSPAN14promoter) was used as control.

10,000 HMC3 cells were plated per well in a 96 well plate format and transfected with 200ng of each pGL4.10-TSPAN14promoter/enhancer or pGL4.10-TSPAN14promoter plasmid, 20ng of the reference plasmid and 145ng of transfection carrier DNA using Lipofectamine 3000 reagent [ThermoFisher Scientific, cat# L3000001) on the next day after seeding. All samples were plated in quadruplicate.

The activity of Firefly luciferase was measured using Dual-Glo luciferase assay system (Promega, cat# E2920) forty-eight hours after transfection. Firefly to Renilla signal ratios were calculated for all samples and compared between different allelic combinations. The experiment was performed in three biological replicates.

### RNA sequencing

Total RNAs were isolated from two clonal lines (##1 and 2) carrying a homozygous deletion in the *TSPAN14* enhancer region and from four control lines expressing the Cas9 protein and either a non-targeting control guide (NTC1, NTC2 and NTC3) or no guide at all (“empty vector” or EV) as described earlier (see RNA extraction). RNA concentrations were determined using Qubit RNA broad range kit (Thermofisher Scientific, Q10211), while RNA integrity (RI) was measured using Agilent RNA 6000 Nano kit (Agilent Technologies, #5067-1512). High RI values, between 9 and 10, were obtained for all samples. A total of 18 RNA libraries (three per sample, with six samples total) were generated using NEBNext Ultra II Directional RNA library prep kit (New England Biolabs, cat# E7760S) and NEBNext Multiplex Oligos (Dual index primers set 1, New England Biolabs, cat# E7600S) for Illumina according to manufacturer’s instructions. The same amounts of total RNA (453ng) were used for making the libraries for all samples. The removal of ribosomal RNA was achieved using the QIASeq FastSelect kit -rRNA HMR kit (QIAgen, cat# 334386). SPRI Select beads (Beckman Coulter, cat# B23317) were used for all bead purification steps. Library quality was confirmed using the Agilent High Sensitivity DNA kit (ThermoFisher Scientific, cat# Q10210). Sequencing was performed on an Illumina NovaSeq 6000 SP flow cell (50bp PE).

### RNA-seq data analysis

The pair-end fastq files were mapped to genome assembly hg19 by STAR (v2.5.2b) independently for each replicate. GencodeV19 annotation was used for gene feature annotation and the raw read count for gene feature was calculated by htseq-count (v0.11.3) with parameter settings -f bam -r pos -s no -t exon -m union. The gene features localized on chrM or annotated as rRNAs were removed from the final sample-by-gene read count matrix. The differential analysis was performed in R (v4.3.1) using R package edgeR. Briefly, the raw counts on genes features were converted to CPM (read Counts Per Million total reads). The gene features with an expression value of more than 0.9 CPM in at least 3 samples were retained for differential analysis. The trimmed mean of M-values (TMM) method was used to calculate normalization scaling factors. Since batch effects due to processing the libraries in three different days were detected by principal component analysis, removeBatchEffects from the R package limma was used to correct count values to adjust for batch effects. Pre- and post-adjustment matrices were used for principal component analysis (prcomp function in the stats R package) and heatmap visualization of clustering of Pearson correlations among pairs of samples (heatmap.2 function in the ggplot R package) to visualize the effects of batch correction. For differential expression analysis, the quasi-likelihood negative binomial generalized log-linear (glmQLFit) approach was applied with model fitting ∼0+condition+batch. The differential expression genes (DEGs) between enhancer KO clones and NT controls were identified with cut-off FDR <0.05 and absolute logFC >0.58.

### Pathway analysis

Statistical enrichment analysis was performed with PANTHER v.19 (accessed 2/21/2025) using all expressed genes (n=16,179) ranked by log2FC * -log10pvalue as the input dataset and “PANTHER GO-Slim Biological Process” as the annotation set. Results were sorted hierarchically and the top, most specific term was retained from each functional class. Terms with an FDR <0.05 were considered significant and were plotted using R packages ggplot2, RcolorBrewer, and patchwork.

### Phalloidin Staining and Quantification

Three NTC control cell lines and two *TSPAN14* enhancer knockout lines were seeded to a 96-well plate at a density of 2,000 cells per well. Three wells were seeded per each condition. On the following day, the cells were fixed using 4% PFA for 10 minutes at room temperature then permeabilized with 0.1% Triton X-100 in PBS for 10 minutes at room temperature. Samples were then blocked using 1% BSA in PBS for 1 hour at room temperature on a shaker. After blocking, samples were stained for 1 hour at room temperature on a shaker using 0.5µL of Phalloidin (Fisher Scientific, cat# A12379) per 200uL of 1% BSA and counterstained with DAPI at a 1:20,000 dilution. Stained samples underwent automated imaging on Zeiss widefield microscope at 10x magnification capturing 9 frames per well.

Quantification of the imaging was performed using CellProfiler and R. Individual nuclei were segmented using DAPI staining. The nuclear segmentation was expanded by 5 pixels and then the nuclear objects were subtracted to create a ring from which the average cytoplasmic fluorescence intensity of phalloidin was calculated in Cell Profiler. In R, the mean fluorescence intensity of all single cells was calculated per well, which was used to perform a two-sample t-test between all NTC versus enhancer KO wells.

### IL-6 and IL-8 ELISA on *TSPAN14* enhancer deletion clones

#### Cell culture media sample collection

Two *TSPAN14* enhancer deletion lines (##1 and 2), two non-targeting guide control lines (NTC1, NTC2) and an empty vector (EV) control line were plated in a 6 well plate format, 100,000 cells per well. Three days after the seeding, cell culture media samples were collected from each well. The volumes of all samples were measured and used to calculate total amounts of interleukins in each sample (see below). The samples were spun down at 1000g for 5min., and the supernatants were aliquoted and stored at −80°C until further processing.

#### ELISA assays

ELISA assays were performed using human IL-6 ELISA (Invitrogen, cat# EH2IL6) and IL-8 ELISA (Invitrogen, cat# KHC0081) kits following manufacturer’s instructions. Briefly, individual aliquots of cell culture media samples were thawed and pre-warmed to room temperature; all other reagents were also equilibrated to room temperature before use. 50µl of each sample were added per well to 96 well ELISA plates in duplicate. IL-6 or IL-8 standards were reconstituted in ultrapure water, and a series of standard curve dilutions was made per manufacturers’ instructions, using HMC3 cell culture media (EMEM, 10% FBS) and a standard diluent, respectively. All procedures were done according to the commercial protocols included in these kits. The absorbance of the final colorimetric products of these ELISA reactions was measured at 450nm using GloMax Discover plate reader (Promega). To determine the total amount of IL-6 and IL-8 in each sample, interleukin concentrations were multiplied by the volumes of respective cell culture media samples. These values were then divided by the total amounts of protein isolated from adherent cells in each well/sample (see below) for normalization.

#### Cell pellet collection from adherent cells, protein extraction and BCA assay

After cell culture media collection, the adherent cells were rinsed in 1xPBS and lifted using TrypLE reagent (Life Technolologies, cat# 1260402). Detached cells were spun down at 130g for 5min., and cell pellets were flash-frozen and stored at −80°C until further processing. Protein lysis and extraction were done in 150µl of the RIPA+ lysis buffer (25mM TRIS-HCl, pH 7.6, 150mM NaCl, 1% NP-40, 1% sodium deoxycholate, 0.1% SDS (ThermoFisher Scientific, cat# 89900)), containing EDTA-free cOmplete Mini protease inhibitor cocktail (1 tablet per 10ml; Millipore-Sigma, cat# 11836170001), PhosStop phosphatase inhibitor cocktail (1 tablet per 10ml; Millipore-Sigma, cat# 05892970001) and 10µ/ul of DNAse I (Millipore-Sigma, cat# 04716728001). Protein lysates were kept on ice for 30min. and then sonicated using a probe sonicator (Fisher Scientific, 40Hz, 2 pulses, 30s each, with a 5sec interval between the pulses). The lysates were spun down at 14000 rpm for 15min. at 4°C, and supernatants were aliquoted and stored at −80°C until further processing. Protein concentrations were determined using the BCA assay based on a standard curve for the bovine serum albumin protein (Thermo Scientific™ Pierce™ BCA Protein Assay Kit, cat# 23225). Cell extracts were diluted to fit the standard curve and loaded on an assay plate in duplicate. Total amounts of protein were determined by multiplying protein concentrations by the total volumes of each protein extract and then used for normalizing the results of the ELISA assay as described above.

---

**Supplementary Table 1.**
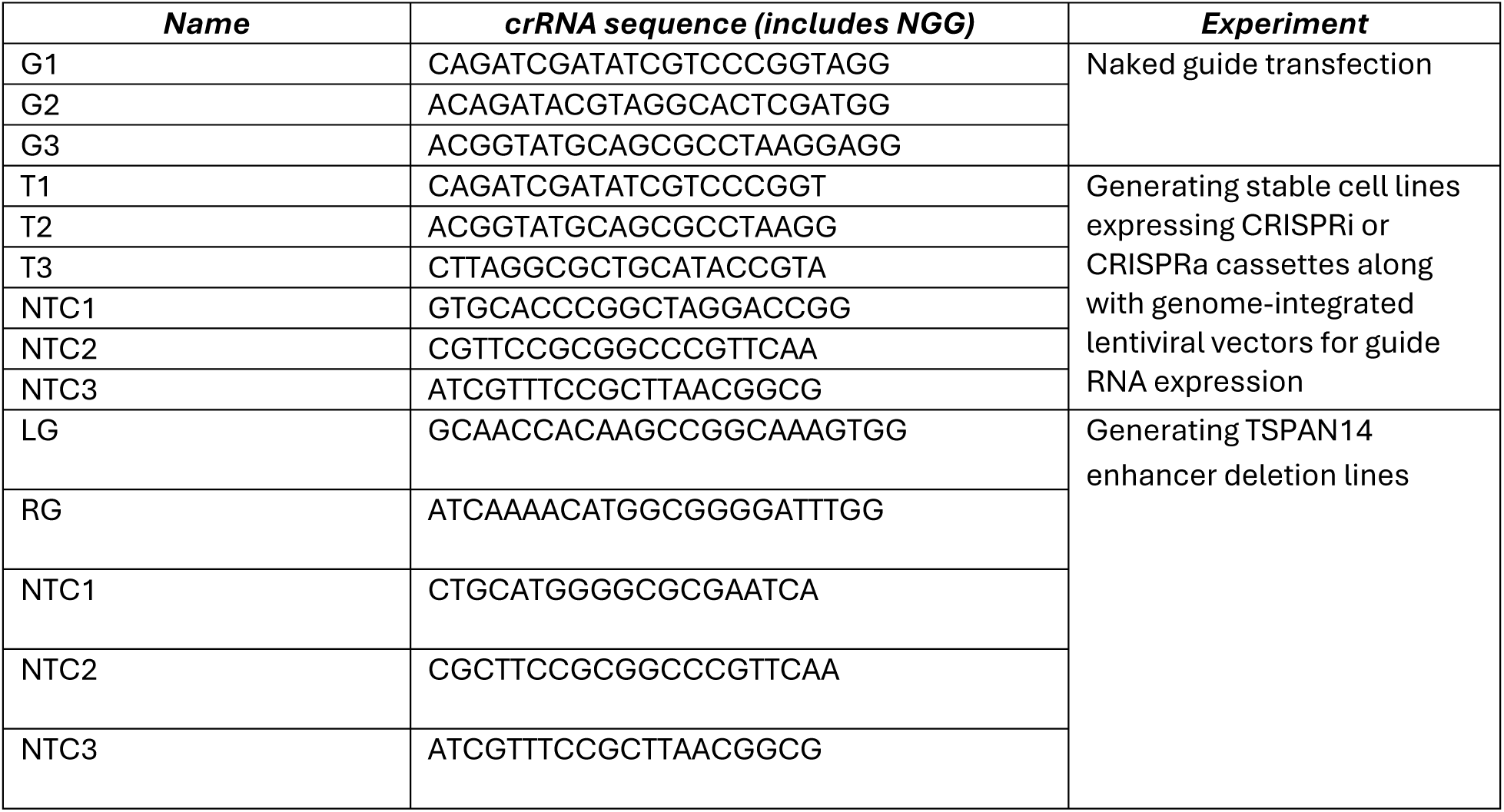
DNA sequences for guide RNAs utilized in CRISPR experiments.

