## Supplementary material for "Integration of Alzheimer’s GWAS, 3D genomics, and single-cell CRISPRi non-coding screen implicates causal variants in a microglial enhancer regulating *TSPAN14*": Suppl. Figure 1

**a)** Transduction efficiency measured by flow cytometry

| | Replicate 1 | | Replicate 2 | | Replicate 3 | |
|---|---|---|---|---|---|---|
| | Count | % Parent | Count | % Parent | Count | % Parent |
| All Events | 20,000 | 100% | 20,000 | 100% | 20,000 | 100% |
| Cells | 15,146 | 75.73% | 13,537 | 67.69% | 12,705 | 63.53% |
| Singlets | 14,167 | 93.54% | 12,384 | 91.48% | 11,508 | 90.58% |
| BFP+ | 1,035 | 7.31% | 953 | 7.70% | 881 | 7.66% |

**b**) Sorting after puromycin selection

| | Post-Selection | |
|---|---|---|
| | Count | % Parent |
| All Events | 20,000 | 100% |
| Cells | 9,571 | 47.66% |
| Singlets | 8,415 | 87.92% |
| BFP+ | 6,051 | 71.91% |

**c)** Dot plots of cells sorted for downstream single-cell RNA sequencing

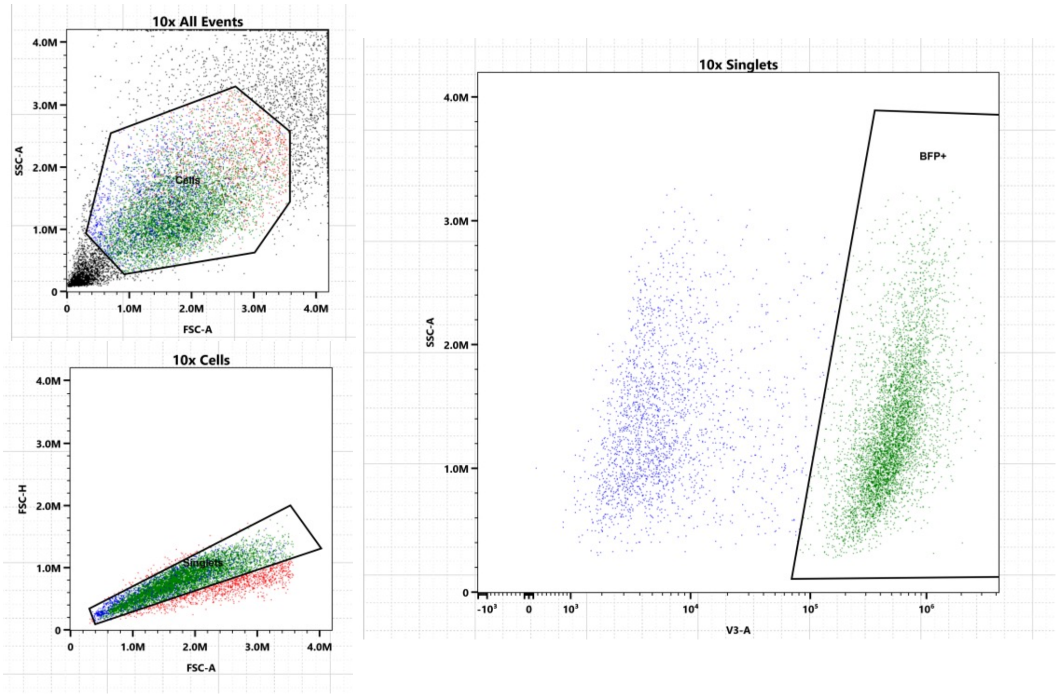


**Supplemental Figure 1. Transduction efficiency of the CRISPRi lentiviral library. a:** Transduction of the sgRNA library (harboring a BFP fluorescent tag) was performed on the HMC3 CRISPRi helper line on control plates alongside the library transduction destined for single-cell RNA sequencing. These cells were not treated with puromycin selection in order to measure the MOI of transduction. All three replicates demonstrated low MOI, where less than 10% of cells were expressing BFP**. b, c:** After 4 days of puromycin selection, remaining cells were selected for BFP expression.
