## Supplementary material for "Integration of Alzheimer’s GWAS, 3D genomics, and single-cell CRISPRi non-coding screen implicates causal variants in a microglial enhancer regulating *TSPAN14*": Suppl. Figure 2

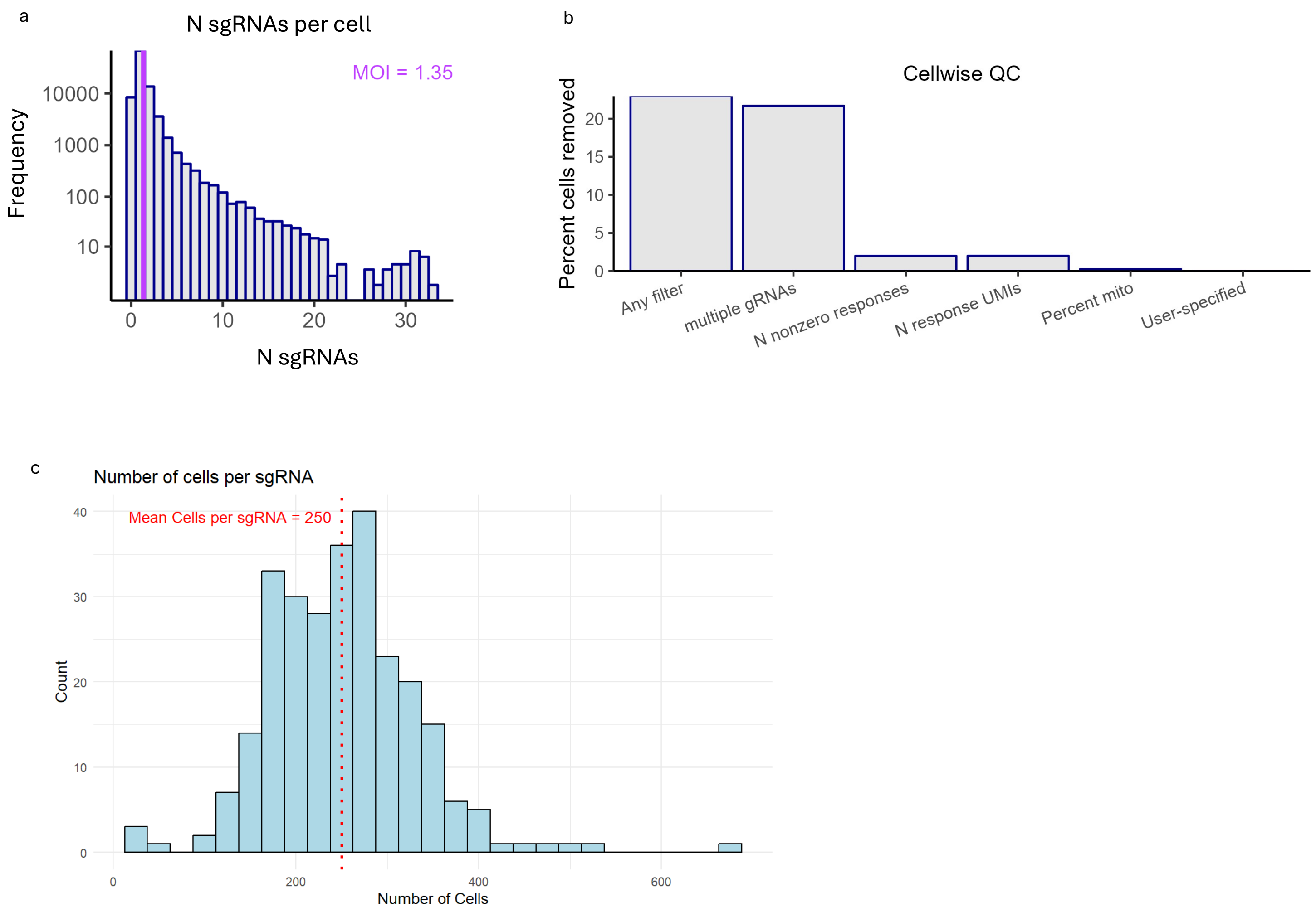


**Supplemental Figure 2. Quality control (QC) of CRISPRi screen. a**: Number of gRNA counts per cell. **b**: Cellwise QC showing percent of total cells removed per respective filter **c**: Number of cells per sgRNA after the gRNA-to-cell assignment step in SCEPTRE. The mean number of cells per guide is shown in red.
