## Supplementary material for "Integration of Alzheimer’s GWAS, 3D genomics, and single-cell CRISPRi non-coding screen implicates causal variants in a microglial enhancer regulating *TSPAN14*": Suppl. Figure 3

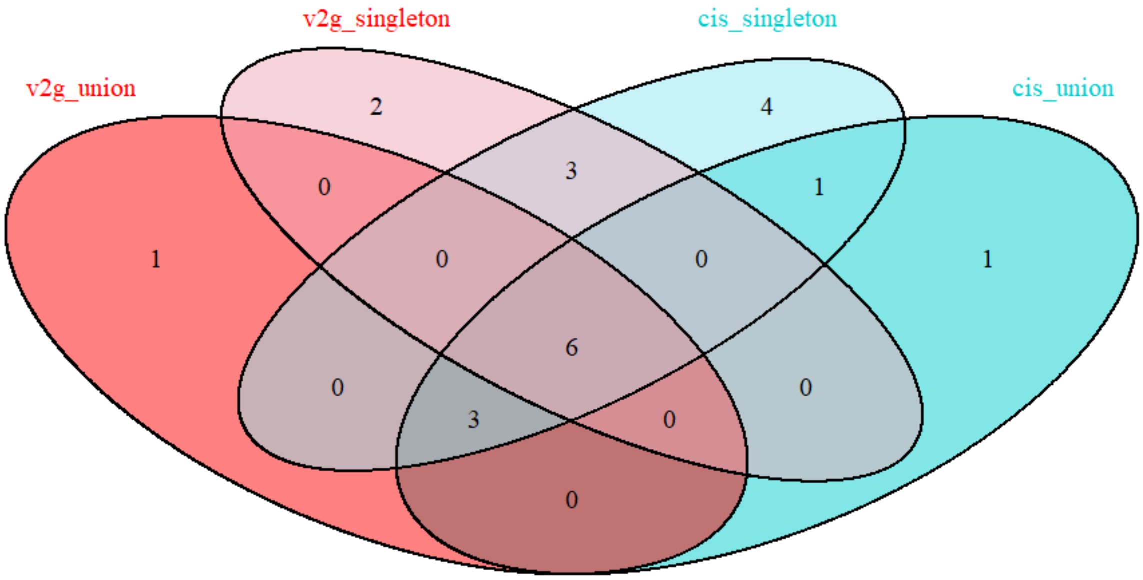


**Supplemental Figure 3. Comparison among hits identified in four different SCEPTRE analyses.**
We analyzed the CRISPRi screen with SCEPTRE using four different approaches. v2g_union: candidate genes from our variant-to-gene mapping approach using "union" parameter for the guides; v2g_singleton: candidate genes from our variant-to-gene mapping approach using "singleton" parameter for the guides; cis_union: all genes included within a 500kb region surrounding the candidate CREs using "union" parameter for the guides; cis_singleton: all genes included within a 500kb region surrounding the candidate CREs using "singleton" parameter for the guides.
