## Supplementary material for "Integration of Alzheimer’s GWAS, 3D genomics, and single-cell CRISPRi non-coding screen implicates causal variants in a microglial enhancer regulating *TSPAN14*": Suppl. Figure 4

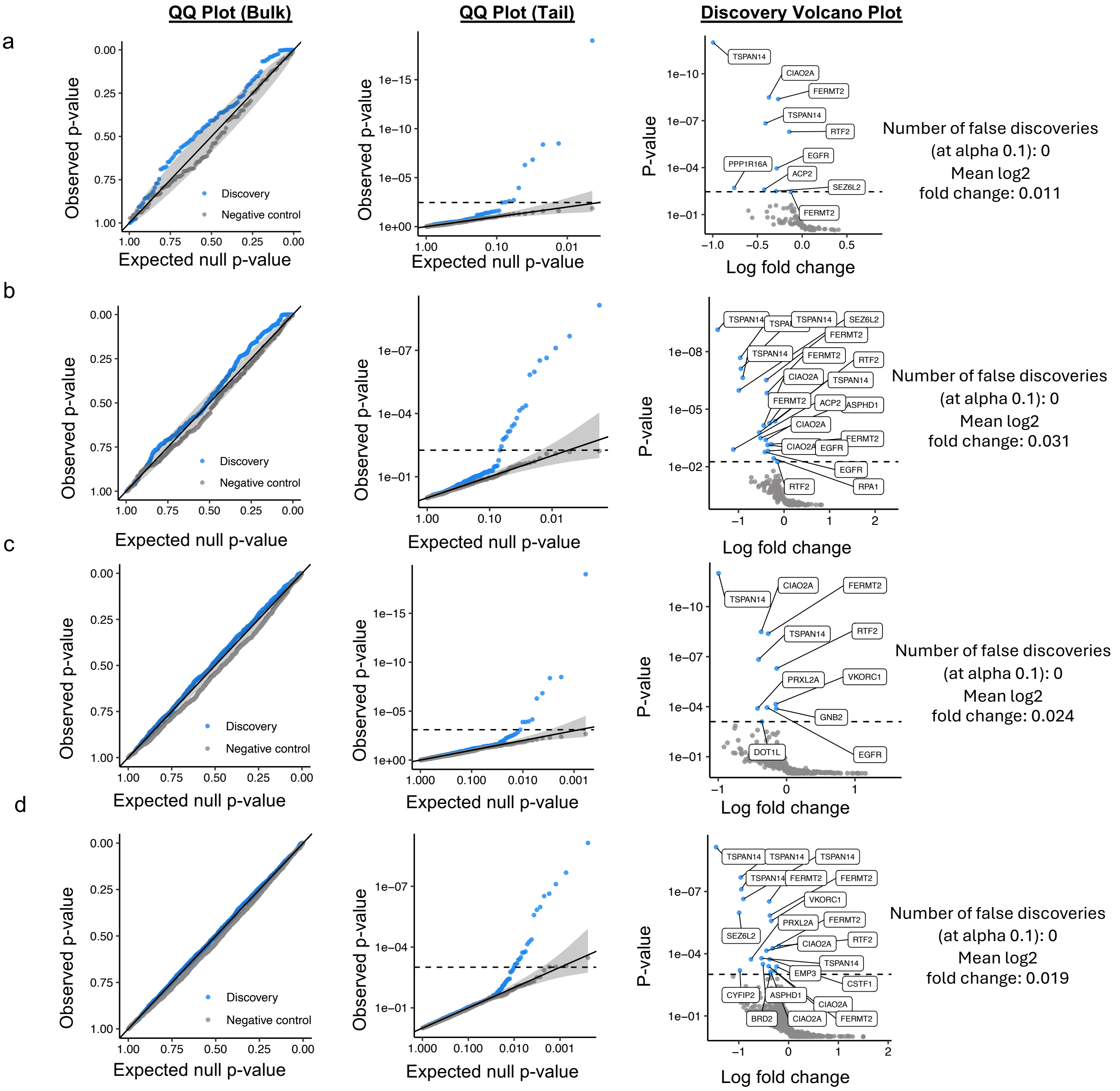


**Supplemental Figure 4. QQ plots (bulk and tail), volcano plots, number of false discoveries found during calibration, and the mean log2 fold change of all pairs for the four different SCEPTRE analyses of the CRISPRi screen. a**: v2g union with candidate genes from our variant-to-gene mapping approach using “union” parameter for the guides; **b**: v2g singleton with candidate genes from our variant-to-gene mapping approach using “singleton” parameter for the guides; **c**: cis union with all genes included within a 500kb region surrounding the candidate CREs using “union” parameter for the guides; **d**: cis singleton with all genes included within a 500kb region surrounding the candidate CREs using “singleton” parameter for the guides.
