## Supplementary material for "Integration of Alzheimer’s GWAS, 3D genomics, and single-cell CRISPRi non-coding screen implicates causal variants in a microglial enhancer regulating *TSPAN14*": Suppl. Figure 5

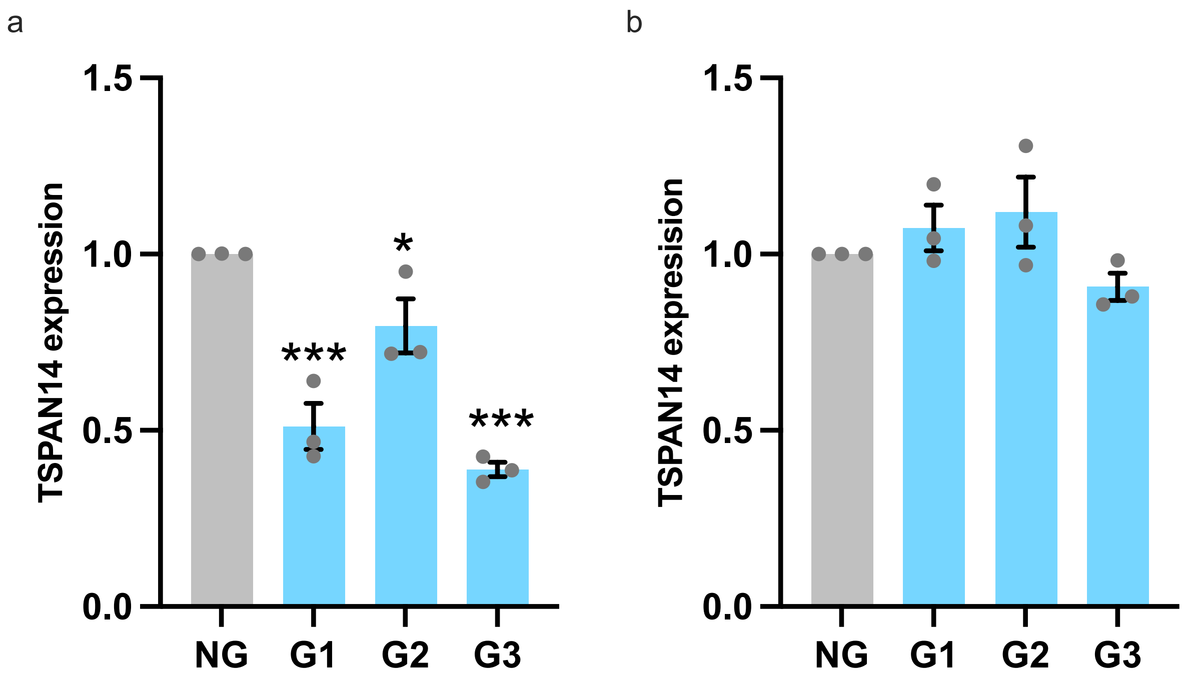


**Supplemental Figure 5: CRISPR interference and activation of the *TSPAN14* intronic region harboring AD-risk variants rs7080009, rs1870138, and rs1870137 in microglial cells using "naked" guides.** Three gRNAs targeting this region were transfected in HMC3 helper lines expressing either dCas9-KRAB (CRISPRi) or dCas9-VP64 (CRISPRa). **a**: All three guides showed a significant downregulation of *TSPAN14* in the CRISPRi line compared to the no guide control. b: no difference in *TSPAN14* expression effect was detected in the CRISPRa experiment. gRNA transfected samples: G1, G2, G3. Mock transfection with no gRNA added: NG. Bar plots show means with SEM error bars; N=3. One-way ANOVA: CRISPRi $P=1.3\times10^{-4}$; CRISPRa *P*=n.s.; pairwise t-test with BH correction: *** $P<5\times10^{-4}$, * $P<0.05$.
