## Supplementary material for "Integration of Alzheimer’s GWAS, 3D genomics, and single-cell CRISPRi non-coding screen implicates causal variants in a microglial enhancer regulating *TSPAN14*": Suppl. Figure 6

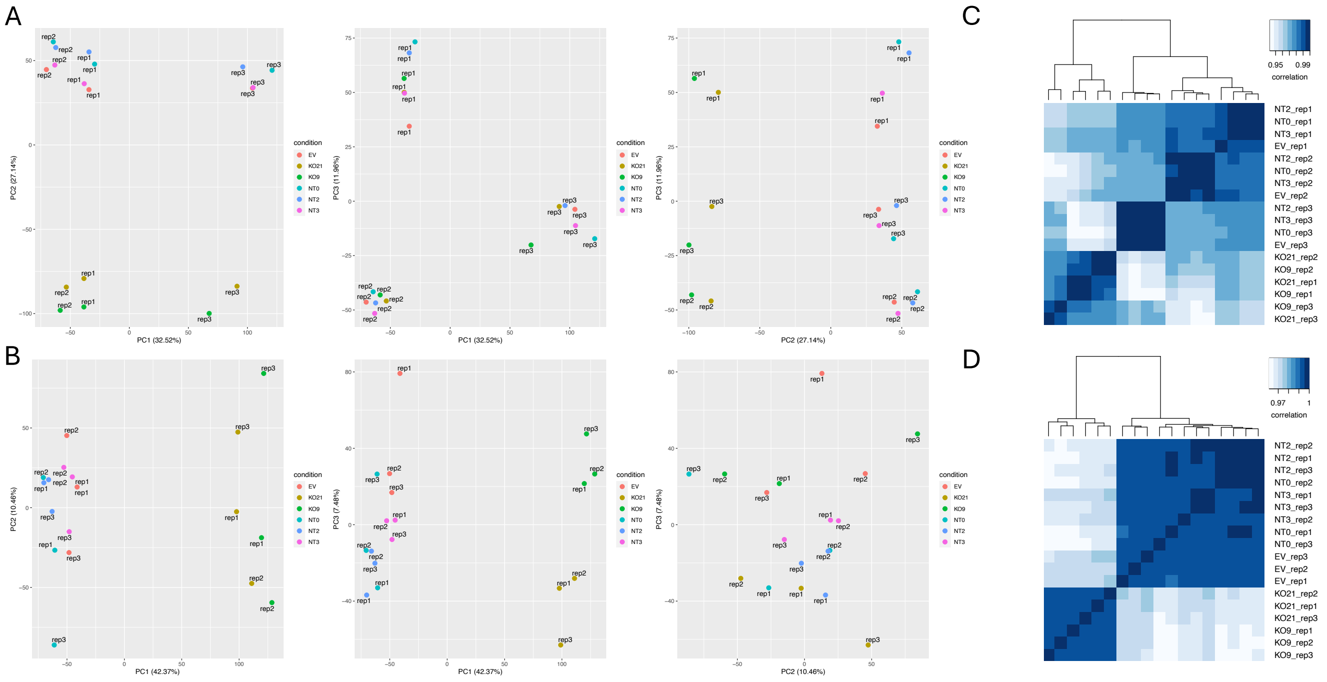


**Supplemental Figure 6. Transcriptomic analysis of TSPAN14 enhancer KO clones: QC and batch effect removal.** RNA-seq was performed on two KO clonal lines (KO9 and KO21), three control clones harboring non-targeting (NT) guides, and one control harboring the empty vector (EV). **A, C**: PCA analysis and heatmap clustering revealed the presence of batch effects (due to the libraries being processed on different days). **B, D**: after adjusting for batch effects, all technical replicates clustered together, with a clear separation by condition (i.e. Mock/EV vs KO).
